# Retrovirus-induced leukemia – hijack of T-cell activation mechanisms revealed by single-cell analysis

**DOI:** 10.1101/2021.09.01.458291

**Authors:** Benjy Jek Yang Tan, Kenji Sugata, Omnia Reda, Misaki Matsuo, Kyosuke Uchiyama, Paola Miyazato, Vincent Hahaut, Makoto Yamagishi, Kaoru Uchimaru, Yutaka Suzuki, Takamasa Ueno, Hitoshi Suzushima, Hiroo Katsuya, Masahito Tokunaga, Yoshikazu Uchiyama, Hideaki Nakamura, Eisaburo Sueoka, Atae Utsunomiya, Masahiro Ono, Yorifumi Satou

## Abstract

Human T-cell leukemia virus type 1 (HTLV-1) mainly infects CD4^+^ T-cells and induces chronic, persistent infection in infected individuals with some progressing to develop adult T-cell leukemia/lymphoma (ATL). Whilst HTLV-1 alters cellular differentiation, activation and survival, it is unknown whether and how these changes contribute to malignant transformation of infected T-cells. In this study, we used single-cell RNA-Seq and TCR-Seq to investigate T-cell differentiation and HTLV-1-mediated transformation processes. We analyzed 87,742 single cells from peripheral blood of 12 infected and 3 uninfected individuals. Using multiple independent bioinformatic methods, we demonstrated that naïve T-cells dynamically change into activated T-cells including infected cells, which seamlessly transitioned into ATL cells characterized by clonally expanded, highly-activated T-cells. Notably, the more activated ATL cells are, the more they acquire Treg signatures. Intriguingly, HLA class II genes were uniquely induced in infected cells, further upregulated in ATL cells and was induced by viral protein Tax. Functional assays revealed that by upregulating HLA class II, HTLV-1-infected cells can act as tolerogenic antigen presenting cells (APCs) to induce anergy of antigen specific T-cells. In conclusion, our study revealed the in vivo mechanisms of HTLV-1-mediated transformation and immune escape at single-cell level.

**Graphical Abstract:** 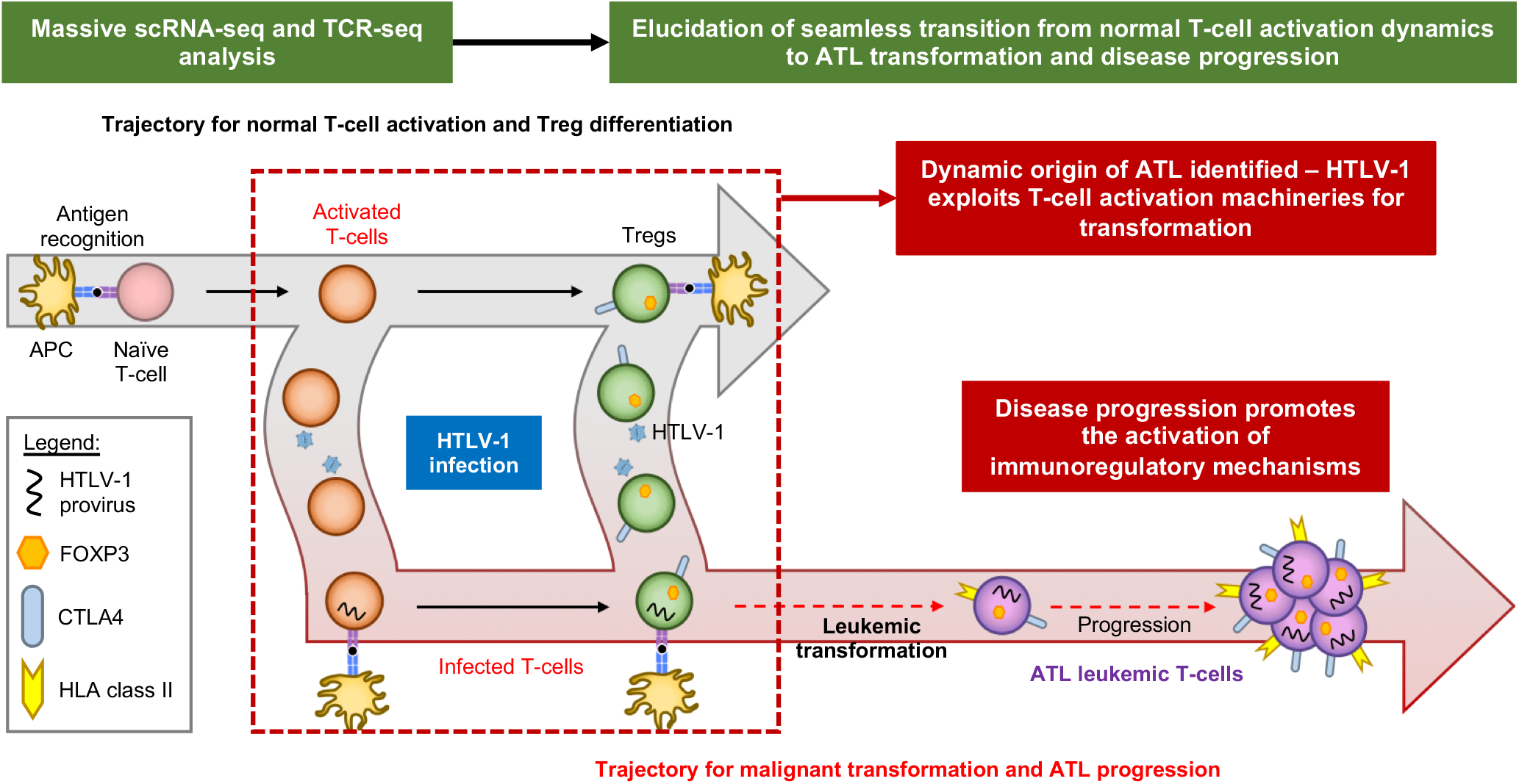

## Introduction

The human T-cell leukemia virus (HTLV-1), the first human retrovirus discovered(1), infects at least 5-10 million people worldwide. HTLV-1 inserts its genome into CD4^+^ T-cells and exists as a provirus in the host cells. After integration, the provirus mainly remains silent and rarely produces viral particles, and the majority of infected individuals remain as asymptomatic carriers (AC) throughout their lifetime(2). However, approximately 5–10% of infected individuals develop CD4^+^ T-cell leukemia/lymphoma, namely adult T-cell leukemia/lymphoma (ATL) decades after infection. Underlying pathogenic mechanisms remain elusive. ATL has two major subtypes: (I) aggressive-type ATL with fast-growing leukemic cells, including acute ATL and lymphoma-type ATL; and (II) indolent-type ATL with slow disease progression, including chronic ATL and smoldering ATL(3). Both of the ATL subtypes have long incubation times, which hampers experimental investigations. Classical studies defined CD25 (IL-2 receptor alpha chain) as a marker for ATL cells(4). *CD25* expression is induced in normal T-cells upon antigenic recognition and subsequent T-cell receptor (TCR) signaling. In addition, regulatory T-cells (Treg), a subset of CD4^+^ T-cells, constitutively express *CD25* and this expression is further induced when they get activated. Recent studies identified other markers for ATL cells, such as FOXP3(5, 6), which is also expressed in Treg and some activated T-cells(7-9). These suggest that ATL transformation involves mechanisms related to either activated T-cells or Treg or both. However, the expression of those markers is variable between and within patients(5, 10). In addition, all those molecules are dynamically regulated in normal T-cells both in steady-states and during T-cell responses at each single T-cell level.

Meanwhile, the viral genes *tax* and *HBZ* have been identified to play key roles in leukemic transformation(11). Tax activates multiple cellular signaling pathways including NF-κB and enhances the survival of host cells(12). HBZ promotes CD4^+^ T-cell hyperproliferation and differentiation towards effector/memory phenotype(13, 14). Intriguingly, activating mutations in ATL genome are accumulated preferentially in genes involved in the NF-κB signaling pathway such as *PLCG1, PRKCB* and *CARD11*, which are a part of T-cell receptor (TCR) signaling*(15)*. However, it is not known how these mutations and viral proteins synergistically and/or cooperatively lead to ATL in its long latent period.

These two lines of evidence support that HTLV-1 infection and subsequent leukemogenesis are closely associated with TCR signaling pathway and CD4^+^ T-cell activation and differentiation. Since these are all dynamically regulated in different normal and infected T-cell populations, single cell-level analysis is required to fully understand how HTLV-1 infection controls normal T-cell mechanisms to transform them into ATL cells. In this study, we cross-analyzed healthy donors, ACs, and ATL patients using single-cell RNA-seq (scRNA-seq) and T-cell receptor sequencing (TCR-seq), aiming to reveal single-cell pathways for in vivo transformation of infected T-cells into leukemic cells.

## Results

### Single-cell transcriptome analysis of PBMCs from HTLV-1-infected individuals

In this study, we focused on indolent-type ATL which shows a relatively slow disease progression. In addition, we included healthy individuals to coherently analyze the whole spectrum of physiological T-cell activation and differentiation in combination with disease progression and oncogenic transformation in ATL patients. We generated droplet-based 5’ scRNA-seq and TCR-seq libraries from PBMCs of 16 individuals comprising of 4 asymptomatic carriers (AC1-AC4), 9 ATL patients (3 smoldering ATL, SML1-SML3; 5 chronic ATL, ATL1-ATL7 and 1 lymphoma-type ATL, ATL4) and 3 healthy donors (HD1-HD3) (Figure 1A and Supplemental Table 1). In total, scRNA-seq profiles from 87,742 cells passed quality filters and included 19,903 cells for AC; 16,357 for SML; 40,119 cells for ATL and 11,363 cells for HD (Supplemental Figure 1A). We identified 25 single-cell clusters based on scRNA-seq profiles, including T-cells, B-cells, NK cells, myeloid cells, megakaryocytes and erythrocytes (Supplemental Figure 1, B-D).

**Figure 1:**
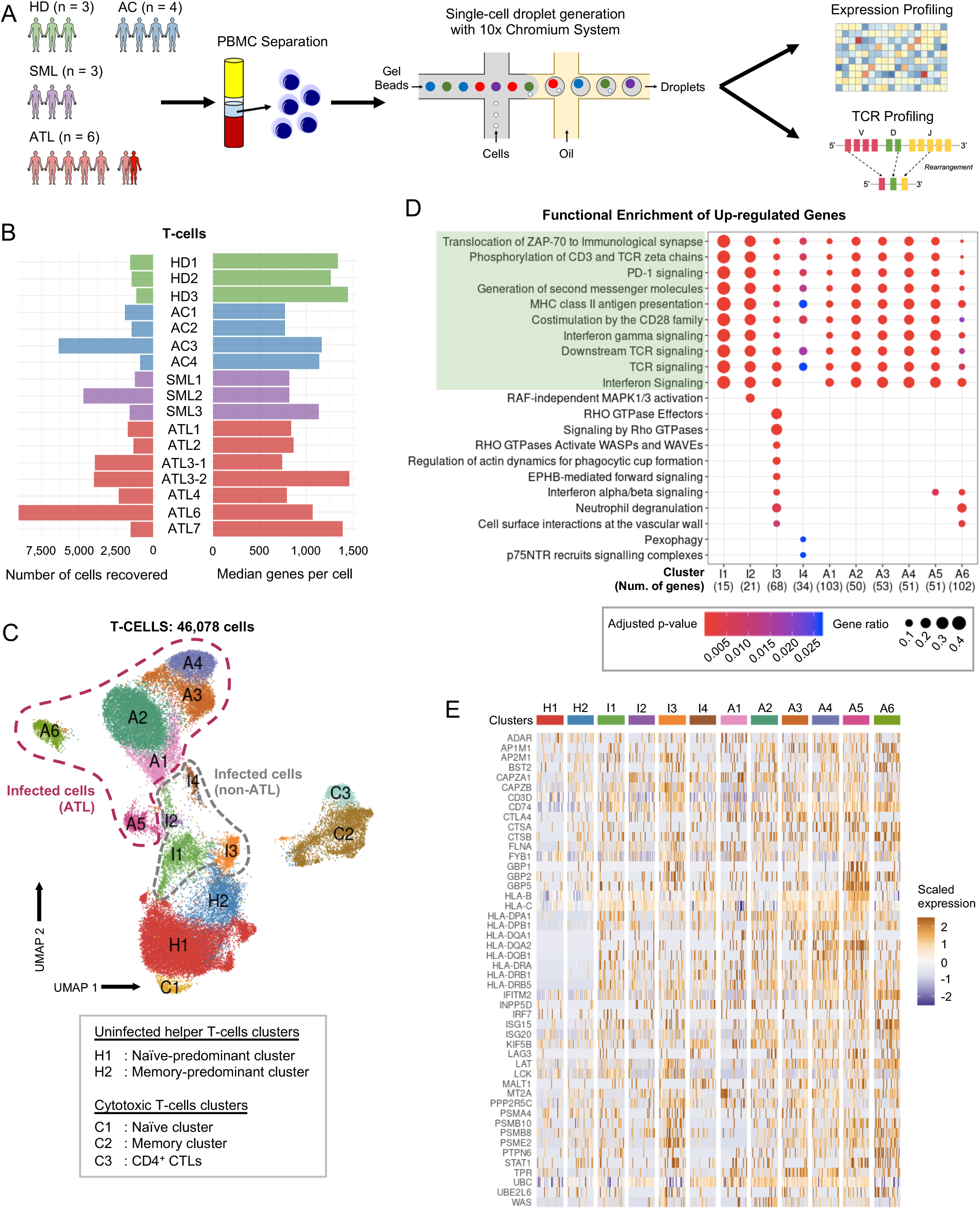
Single-cell transcriptional profiling of HTLV-1-infected individuals. (**A**) Schematic representation of experimental workflow. (**B**) The number of T-cells recovered that passed quality controls (left) and their median number of genes per cell (right) for each of the individuals (3 healthy donors, HD1-HD3; 4 asymptomatic carriers, AC1-AC4; 3 smoldering ATL, SML1-SML3; and 7 ATL, ATL1-ATL7). (**C**) 2-D UMAP visualization showing the 4 different groups of T-cells which we identified. (**D**) Dot plot shows the shared and distinct Reactome pathways of upregulated genes for each infected cluster. Common pathways between all clusters are highlighted in green. (**E**) Heatmap shows the expression of the genes making up the common pathways highlighted in green in panel **D** for all clusters.

Next, we performed in silico sorting of T-cells and clustering analysis in order to understand the dynamics of T-cell differentiation and ATL transformation at the single-cell level. In total, we obtained 46,078 T-cells which includes 10,576 from AC; 7,514 from SML; 23,845 from ATL and 4,143 from HD (Figure 1B). We identified 15 distinct T-cell clusters (Figure 1C). Interestingly, cells from HDs, ACs and SMLs were mixed altogether in different clusters (H1-H2 and C1-C3) whereas cells from ATLs formed distinct clusters on their own (Supplemental Figure 2, A and B). Three CD8^+^ cytotoxic T-cells clusters (C1-C3) were identified while the remaining clusters were classified as CD4^+^ helper T-cells based on their key gene expression (Supplemental Figure 2C). We further confirmed our annotations by mapping our data to a reference dataset (Supplemental Figure 3, A and B). Cells contributing to the clusters I1-I4 and A1-A6 were identified as HTLV-1-infected cells (*HTLV-sense*^+^ *HBZ*^+^ *CADM1*^+^ *CCR4*^+^ *CD40LG*^lo/−^ *CD7*^lo/−^ *DPP4*^lo/−^) (Supplemental Figure 2D)(16, 17). To confirm our annotation for HTLV-1-infected cells, we measured and compared the proviral load between CADM1^−^ CD7^+^ cells and CADM1^+^ CD7^−/+^ cells, and revealed that CADM1^+^ CD7^−/+^ cells are HTLV-1-infected as evidenced by the high proviral loads (>60%) (Supplemental Figure 4C). As ATL cells are known to be highly expanded T-cell clones, we analyzed paired single-cell TCR sequences to identify clonally expanded cells. We noted that among all clusters derived from ATLs, several primarily consisted of cells with a single, highly expanded clone (clusters A1-A6) (Supplemental Figure 4A). Intriguingly, some cells from healthy individuals clustered together with HTLV-1-infected cells from multiple ATL patients (Supplemental Figure 2A and 4B). This suggests that some of the same T-cell clones as ATL cells exhibit considerably similar transcriptional profile to normal T-cells in healthy individuals.

Next, we performed a differential expression analysis in order to identify shared characteristics between all infected/ATL clusters. Pathway analysis of upregulated genes showed that these clusters were significantly enriched in genes used for 10 TCR- and activation-related pathways including TCR signaling, antigen presentation and immune checkpoints (Figure 1D). To identify the key genes accounting for the TCR-related pathways, we looked into the expression profile of all genes in the 10 shared pathways and found that there was a strong enrichment of human leukocyte antigen (HLA) class II genes among infected/ATL clusters (Figure 1E). A similar analysis of downregulated genes showed neither common pathways nor enriched genes among these clusters (Supplemental Figure 5, A and B).

### ATL cells highly express co-stimulatory receptors and genes downstream of TCR signaling

While most of ATL cells highly express *IL2RA* (CD25), a key feature for activated T-cells and Treg, they may show variable expression of genes used in activated T-cells and Treg markers. We hypothesized that ATL cells in each patient are heterogenous, showing a broad spectrum of the gene expression profiles of activated T-cells and Treg. In order to test this hypothesis, we applied to the dataset of CD4^+^ T-cells canonical correspondence analysis (CCA), which is a multidimensional method to quantitatively analyze gene expression profiles for the differentiation or activation status of individual single cells in a data-oriented manner(18). We first analyzed the CD4^+^ T-cell data using the reference RNA-seq data for activated T-cells in order to evaluate ‘T-cell activation score’ of individual ATL and non-ATL cells. As expected, the majority of CD4^+^ T-cells from HDs had low T-cell activation scores, which is compatible with their naïve status, while T-cells from HTLV-1-infected individuals had higher scores with ATLs having the highest scores (Figure 2A, left panel and Supplemental Figure 6A, top row). We next used a reference RNA-seq data for Treg and analyzed ‘Treg score’ of individual ATL and non-ATL cells, finding a similar trend; T-cells from HDs show low Treg scores while cells from SML and ATL have remarkably high Treg scores (Figure 2A, right panel and Supplemental Figure 6A, bottom row). Two-dimensional plot of the two CCA scores shows that while HDs harbor T-cells with activated phenotype, infected T-cells and ATL cells spontaneously become activated and acquire Treg phenotype and subsequently progress into extremely activated, which is maintained throughout the ATL phase (Figure 2B and Supplemental Figure 6B). This interaction of transcriptional activities for Treg and activated T-cells was further confirmed by a 2D-CCA analysis which used both of the variables for T-cell activation and Treg. Most ATL cells are found in the quadrant between the T-cell activation and Treg axes, indicating that they are both highly activated and acquire Treg phenotype (Supplemental Figure 6D).

**Figure 2:**
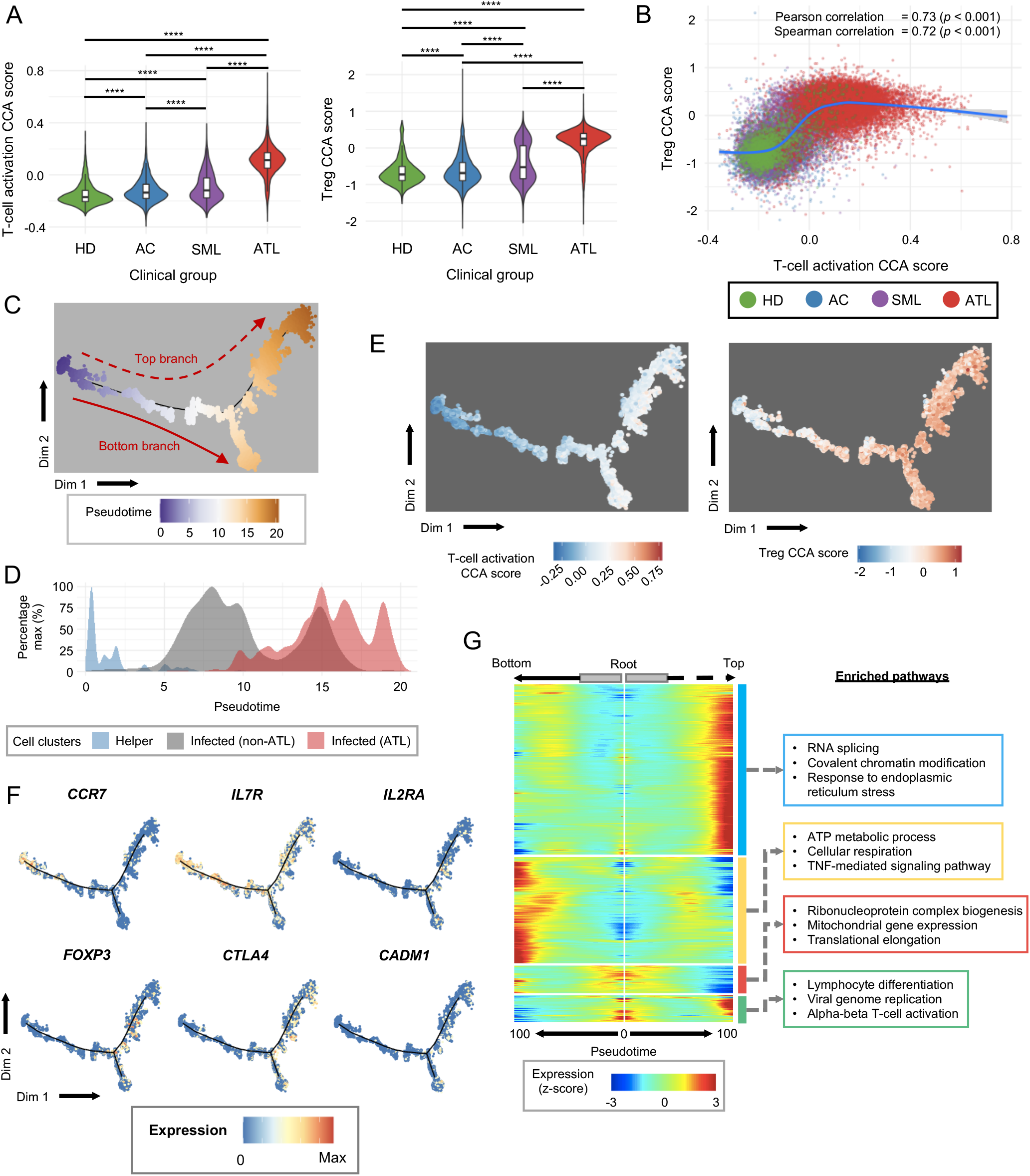
CCA analysis shows that ATL cells acquire Treg phenotype and are highly activated. (**A**) Violin plots show the 1-D CCA scores for T-cell activation and Treg grouped by clinical diagnosis. Box plots in each violin summarize the median (midline) and IQRs. (**B**) Scatter plot shows the correlation between the 1-D CCA score for T-cell activation and Treg. The blue line shows the regression model of the CCA scores for all cells. (**C**) Plot showing the pseudotime trajectory for the entire CD4^+^ T-cells population colored by pseudotime. (**D**) Plot shows the distribution of uninfected helper cells, infected non-ATL cells and infected ATL cells along the pseudotime axis. (**E**) Plot showing the distribution of 1-D CCA scores for T-cell activation and Treg in the pseudotime space in panel **C**. (**F**) Expression of T-cell-related marker genes along the pseudotime axis in panel **C**. (**G**) Split heatmap shows the expression profile of genes which varies as a function of pseudotime and are branch dependent. Pseudotime trajectory begins from the middle of the heatmap (grey box) and moves to the left for the bottom branch and to the right for the top branch. The start of the arrow indicates the bifurcation point in the trajectory in panel **C**. Hierarchical clustering groups the genes into 4 clusters indicated with color boxes on the right with their top 3 enriched pathways indicated. **** p-value < 0.0001 by Wilcoxon rank sum test (**A**).

Additionally, we analyzed the expression of co-stimulatory and co-inhibitory molecules (i.e. immune checkpoints), defining the activation and exhaustion signatures, respectively. In general, HTLV-1-infected cells had higher activation and exhaustion signatures compared to non-infected and normal T-cells (Supplemental Figure 5C). Among the infected cells, the two signatures were the most pronounced in ATL cells (Supplemental Figure 5C). Interestingly, the activation signature was highly variable between HTLV-1-infected and ATL clusters and none of the genes was consistently expressed by all the infected and ATL clusters. On the other hand, most of the HTLV-1-infected and ATL clusters expressed similar exhaustion signature genes including *TIGIT, CTLA4* and *LAG3* (Supplemental Figure 5D). This indicates that infected/ATL cells have a broad spectrum of activation states while harboring a phenotype of chronically stimulated, exhausted T-cells.

We earlier postulated that by using cells from patients diagnosed with indolent ATL, we would be able to capture cells from different stages of oncogenic transformation and disease progression. To address this hypothesis, we applied a pseudotime analysis to the entire CD4^+^ T-cell population, regardless of the sample identity. Pseudotime analysis showed that uninfected helper T-cells clustered in early pseudotime while infected/ATL cells clustered around the middle and end of the pseudotime trajectory (Figure 2, C and D and Supplemental Figure 7A). Interestingly, the trajectory of the cells bifurcated into 2 different branches suggesting that there were two phenotypically different ATL cells. We then assessed how gene expression is regulated across the pseudotime axis using the CCA T-cell activation and Treg scores. As expected, both of the CCA scores gradually and progressively increased along the trajectory, indicating that the transformation of healthy T-cells into ATL cells involves the activation of the infected cells and acquisition of Treg phenotype (Figure 2E). Next, we analyzed the changes in gene expression varying across pseudotime. As expected, T-cells upregulated HTLV-1 infection-related genes such as *CADM1* in the latter half of pseudotime axis. Intriguingly, T-cells from healthy individuals upregulated activation and Treg-related markers including *IL2RA* and *FOXP3* along the pseudotime axis which was seamlessly followed by HTLV-1-infected cells and ATL cells towards the end of the pseudotime axis. On the other hand, markers of naïve cells such as *CCR7* and *IL7R* were downregulated with pseudotime throughout the healthy, infected and ATL cells (Figure 2F). These collectively indicate that our inferred pseudotime trajectory had successfully captured the normal T-cell developmental/differentiation and the HTLV-1-mediated T-cell transformation processes as a continuous physiological and pathological pathway.

### Pseudotime analysis reveals that Treg associated genes such as *IL2RA, FOXP3* and *CTLA4*, and HLA class II genes are upregulated along ATL trajectory

We showed earlier using a collective analysis of all CD4^+^ T-cells, we observed that there were at least 2 phenotypically different ATL cells (Figure 2C). We performed differential expression analysis to reveal the difference between these two branches. Pathway analysis showed that cells in the top branch upregulates genes related to transcription and endoplasmic reticulum stress while genes upregulated in cells of the bottom branch are involved in metabolic process and TNF signaling (Figure 2G). However, as we showed in Figure 1C, the ATL cells mainly formed clusters on their own with some overlaps, indicating that there is a large inter-individual variability of ATL cells. This large inter-individual variations of ATL cells may obscure unique dynamics of ATL cell transformation in each individual. Accordingly, we performed pseudotime analysis by combining cells from healthy donors (n=3) and individual ATL patients, producing a total of 5 pseudotime datasets (Figure 3A). All the 5 datasets successfully identified the trajectory of ATL cells: healthy donors mainly clustered at earlier pseudotimes while cells from ATL clustered in the middle and end (Figure 3B). Expression kinetics of T-cell-related genes showed that along the pseudotime trajectory, expression of naïve-related genes such as *CCR7* was downregulated while HTLV-1 infection-related genes (*CADM1*), T-cell activation and Treg-related genes (*IL2RA, FOXP3, CTLA4*) were upregulated (Figure 3C and Supplemental Figure 8). This shows that the earlier segment of the pseudotime recapitulates the developmental/differentiation pathway of normal CD4^+^ T-cells as they transition from naïve to effector cells. This was seamlessly followed by the later segment which displays the progression of T-cells after HTLV-1 infection in which cells transition from infected, polyclonal cells to highly expanded, monoclonal cells (Figure 3, B and C).

**Figure 3:**
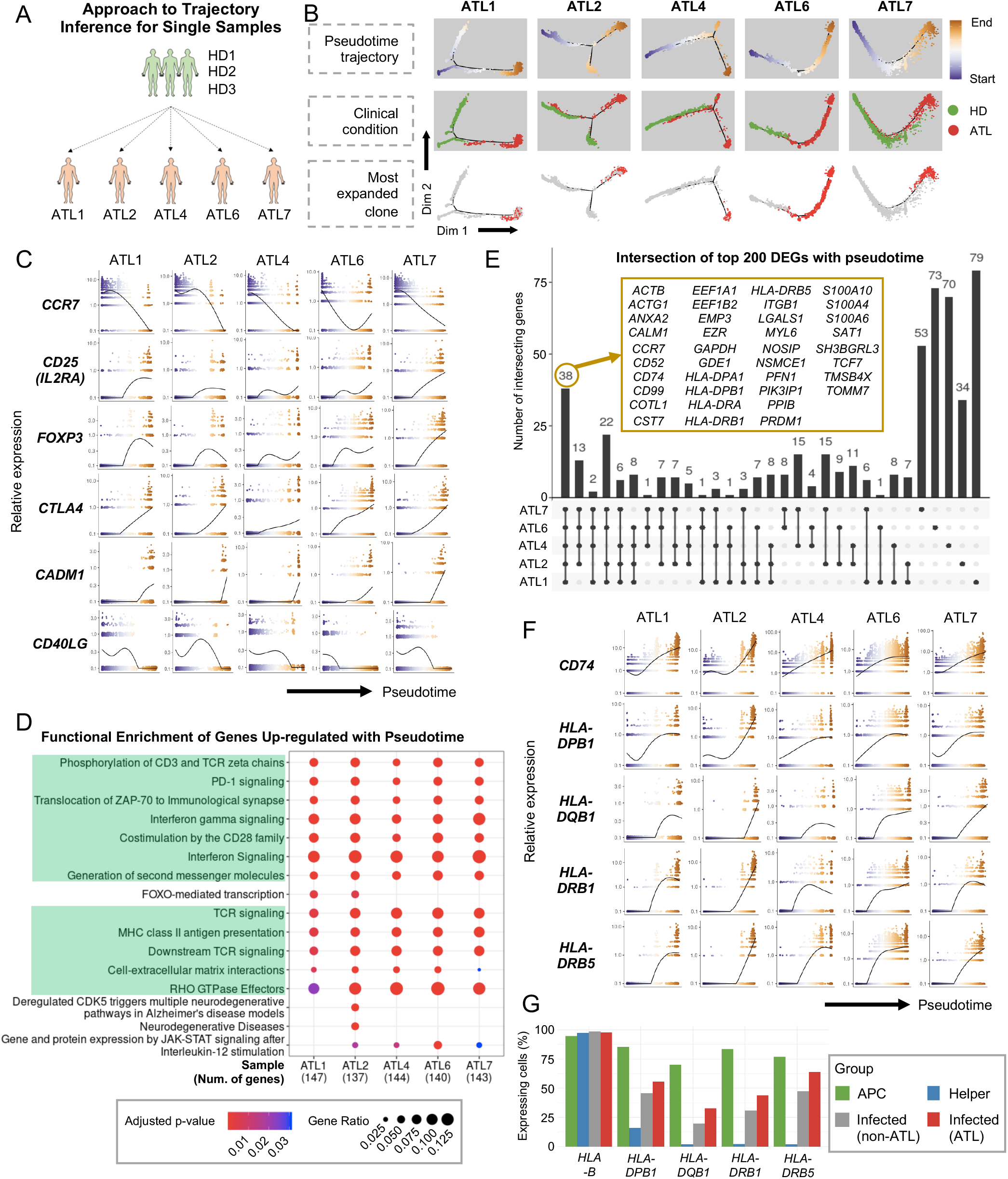
Pseudotime analysis of single ATL samples. (**A**) Schematic figure showing pseudotime analysis is performed using 3 HD and individual ATL cases. (**B**) Plots showing the clusters and pseudotime trajectories for each individual ATL with 3 HD; the first row colors the clusters by pseudotime, the second row colors the clusters by clinical condition and the last row marks the distribution of the most expanded clone for each ATL patient. (**C**) Expression dynamics of T-cell-related genes along the pseudotime axis. Color of the dots represent the cells position along the pseudotime axis as in panel **B**. (**D**) Dot plot shows the shared and distinct Reactome pathways of the top 200 genes varying as a function of pseudotime for each ATL sample. (**E**) Upset plot showing the intersection of the top 200 genes varying as a function of pseudotime for each ATL sample. Genes common to all ATL cases are shown in the yellow box. (**F**) Expression dynamics of HLA class II and related genes along the pseudotime axis. Color of the dots represent the cells position along the pseudotime axis as in panel **B**. (**G**) Bar plot shows the percentage of cells in APCs (B cells and monocytes), uninfected helper T-cells, infected non-ATL cells and infected ATL cells that express HLA class II genes. *HLA-B* is shown here as control.

In order to identify the common factors contributing to ATL progression, we examined the top 200 genes varying as a function of pseudotime for each dataset. We found that the top 200 genes shared some common enriched pathways such as the involvement of interferon and TCR signaling, and antigen presentation by HLA class II. Among the top 200 genes, 36 of them were common for all ATL patients and included several HLA class II genes (Figure 3, D and E). The expression of these HLA class II genes increased along the pseudotime axis, indicating that as cells transformed into ATL cells, they upregulated HLA class II (Figure 3F and Supplemental Figure 8).

HLA class II are mainly found on professional antigen presenting cells (APCs) such as monocytes or B-cells but they can also be upregulated in T-cells upon stimulation. To investigate the significance of this upregulation, we compared the expression level of HLA class II between professional APCs and the various T-cell clusters. We observed that very few uninfected T-cells expressed HLA class II, and that the percentage of HLA class II positive cells increased in HTLV-1-infected cells and was even higher in ATL cells (Figure 3G). We also found that the expression of genes related to HLA class II signaling such as *CIITA, RFX5* and *CD74* are increased in infected/ATL cells in comparison to uninfected cells (Supplemental Figure 10A)(19, 20).

To address if the pseudotime analysis above recapitulated the real time-course of disease progression, we next used a pair of longitudinal samples to perform trajectory inference together with samples from healthy individuals (Figure 4A). Pseudotime analysis showed that the cells from the first time point distributed around the middle of the pseudotime axis, while cells from the second time point mainly clustered towards the end of the pseudotime trajectory (Figure 4B), indicating that ATL cells at the second time point had a more extreme activated phenotype (Supplemental Figure 9A). RNA velocity analysis also showed the similar pattern with cells transitioning from the naïve and memory state into infected cells, further supporting the results by pseudotime analysis (Figure 4C). Notably, cells in both branches exhibited similar gene expression dynamics similar to the single time-point samples in Figure 3C, including the downregulation of *CCR7* and the upregulation of *CADM1* and *FOXP3* across the pseudotime axis (Supplemental Figure 9D). The expression of HLA class II and its related genes was also remarkably induced across the pseudotime axis (Figure 4D), confirming that the upregulation of HLA class II molecules was a key feature of ATL progression in the longitudinal analysis as well.

**Figure 4:**
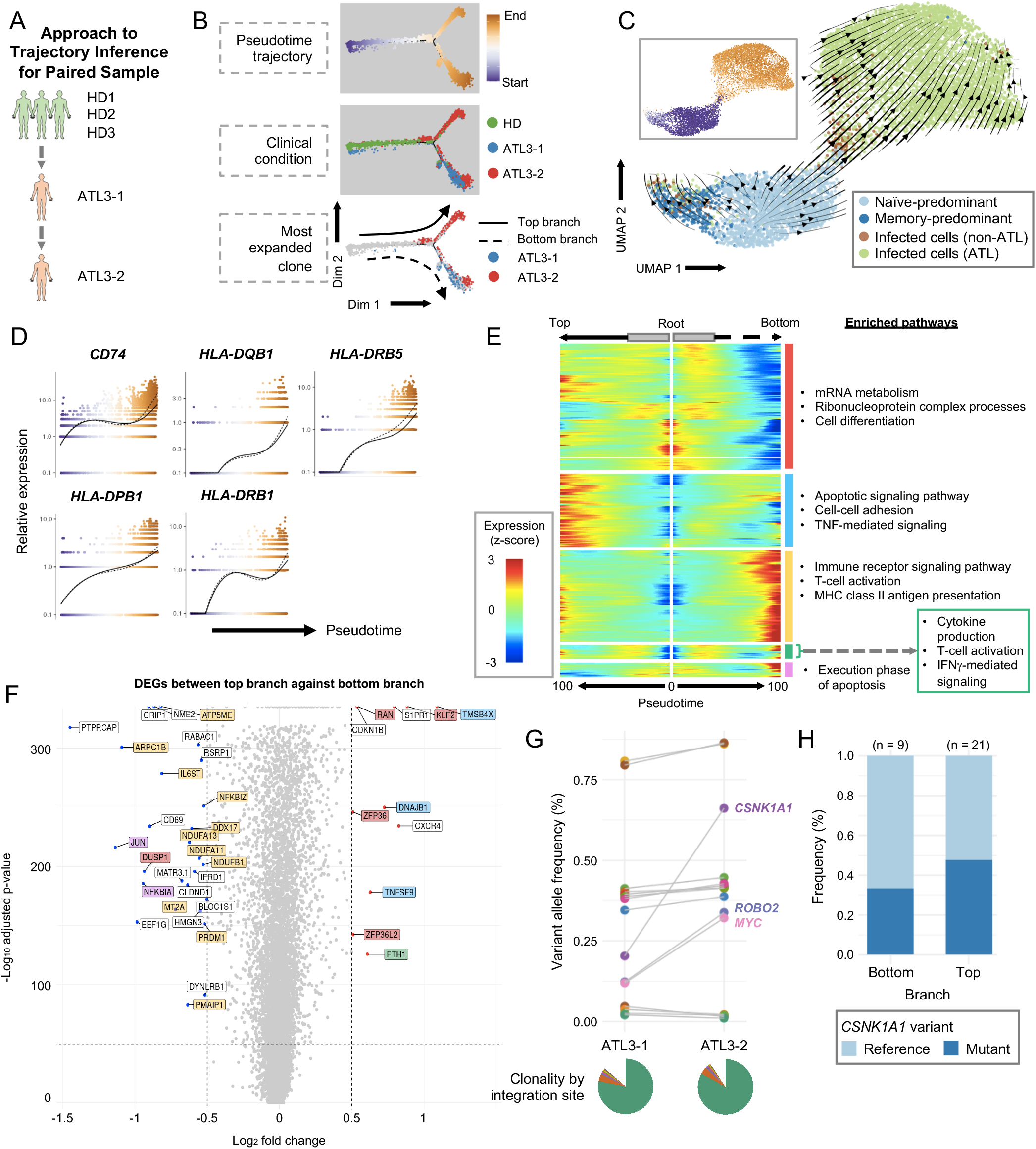
Pseudotime analysis of paired ATL sample. (**A**) Schematic figure showing pseudotime analysis is performed using 3 HD and one paired ATL case. (**B**) Plot showing the clusters and pseudotime trajectory; from top to bottom, clusters are colored by pseudotime, clinical condition and most expanded clone for each ATL. (**C**) Plot shows the RNA velocity in Seurat-identified clusters. The inset box is colored by pseudotime as shown in panel **B**. (**D**) Expression dynamics of HLA class II and related genes along the pseudotime axis. Color of the dots represent the cells position along the pseudotime axis as in panel **B**. (**E**) Split heatmap shows the expression profile of genes which varies as a function of pseudotime and are branch dependent. Pseudotime trajectory begins from the middle of the heatmap (grey box) and moves to the left for the top branch and to the right for the bottom branch. The start of the arrow indicates the bifurcation point in the trajectory in panel **B**. Hierarchical clustering groups the genes into 5 clusters indicated with color boxes on the right with their enriched pathways indicated. (**F**) Volcano plot shows the genes which are differentially expressed between the top and bottom branch. The genes are colored according to the clusters they belong to in the split heatmap in panel **E**. (**G**) Dot and line graph shows the change in frequencies of variant alleles across the two timepoints in the CADM1^+^ CD7^-/+^ population from the paired ATL case as detected using targeted exome-seq. The genes which showed huge changes in frequency are labelled here. The pie chart at the bottom shows the ATL cells clonality identified by HTLV-1 integration site. (**H**) Bar graph shows the frequency of the *CSNK1A1* variant by trajectory branch detected in the scRNA-seq data.

Interestingly, the trajectory of the cells bifurcated into 2 different branches towards the end of the pseudotime axis, indicating that ATL cells in these two branches in the patient are considerably different (Figure 4B). While the ATL cells in both of the branches showed the key activation features as written above, pathway analysis showed intriguing differences between the two branches: T-cells in the top branch significantly used genes for pathways related to TNF-mediated signaling (*TMSB4X*) and cellular adhesion (*TNFSF9*), while T-cells in the bottom branch significantly used genes for T-cell activation (*IL6ST*) and immune signaling (*NFKBIZ*) (Figure 4, E and F).

Next, we addressed if the transcriptome difference between the two branches was due to genomic changes that produced two distinct malignant clones in the patient. Firstly, we examined the distribution of the most expanded clone by TCR-seq for each time point and observed that the clones were similarly distributed between the two branches (Figure 4B, bottom panel and Supplemental Figure 4A), indicating that the ATL cells in the two branches originated from the same TCR clone. To explore the origin of this branching, we first analyzed the genomic landscape of these samples by sorting and analyzing HTLV-1-infected cells (CADM1^+^ CD7^−/+^) from the samples. We found that for both timepoints, there was only one significantly expanded clone based on HTLV-1 integration site distribution, indicating that the ATL cells originated from one parent clone (Figure 4G, pie charts). Next, we performed targeted exome-seq on the sorted cells and identified the somatic mutations present in ATL cells and identified those that differed between the 2 time points (Figure 4G). Subsequently, we analyzed the frequency of these variant alleles in the scRNA-seq data. However, no mutation was found enriched between the two branches from our analysis (Figure 4H), suggesting that the difference between the cells in the two branches is due to either or both of epigenetic modification and/or their reactivities to some signaling (e.g. cytokine milieu) or metabolic status(21, 22). It remains to be elucidated by future studies whether the two major bifurcated fates have any biological and/or clinical significance.

### The expression of HLA class II genes is associated with HTLV-1 infection and the viral protein Tax

We next addressed whether HLA class II expression is associated with HTLV-1 infection. We first examined the expression of HLA class II in HTLV-1-associated cell lines by flow cytometry. We observed that HLA class II expression was upregulated in all HTLV-1-associated cell lines we analyzed but not in HTLV-1-negative ones with the exception of Kit225 cells (Figure 5A). Next, we analyzed freshly isolated CD4^+^ T-cells from PBMC of HTLV-1-infected individuals by comparing HLA class II expression between the uninfected cell fraction (CADM1^−^ CD7^+^) and the HTLV-1-infected / ATL cell fraction (CADM1^+^ CD7^−/+^). We found that the HTLV-1-infected / ATL cell fraction showed a significantly higher level of HLA class II expression compared to the uninfected cell fraction. However, when compared to professional APCs (CD14^+^ monocytes), the upregulation was moderate in HTLV-1-infected and ATL cells (Figure 5, B and C). It is well-established that the HTLV-1 viral genes *tax* and *HBZ* have oncogenic functions. Therefore, to investigate which of these two genes could contribute to the upregulation of HLA class II, we transfected Kit225 cells with expression plasmids containing either HTLV-1 *tax, HBZ* or HIV-1 *nef* as a control. Cells transfected with Tax upregulated HLA class II but no further induction was seen in cells transfected with HBZ or nef (Figure 5D). Level of *tax* expression in fresh PBMC is low but can be induced in bursts and the expression can be induced by ex vivo cultivation(23-25). Thus, to investigate the dynamics of Tax and HLA class II expression, we performed scRNA-seq on PBMCs from 3 infected individuals before and after ex vivo cultivation. Generally, for all 3 samples, clustering analysis of CD4^+^ T-cells revealed 3 clusters consisting of uninfected helper cells, infected cells and a cluster consisting of cells with high expression of HTLV sense strand. Pseudotime analysis inferred 2 trajectories in all the three samples. The first trajectory transitioned from uninfected cells to infected cells while the second trajectory progressed from uninfected to infected and to cells displaying a strong induction of HTLV sense strand transcription (Figure 5E). We next focused on the second trajectory in order to investigate the dynamics of HTLV sense strand transcription and HLA class II expression. We observed that along the trajectory, Tax expression was remarkably induced while the expression of HBZ was variable. The expression of HLA class II also increased along the trajectory and the upregulation occurred concurrently with the increase in Tax expression (Figure 5F). We then aimed to ascertain the mechanism of how Tax leads to the upregulation of HLA class II. We earlier showed that the expression of genes related to HLA class II signaling was also increased in ATL (Supplemental Figure 10A) and one of the genes, *CIITA*, is known to be a master regulator of HLA class II gene expression. We performed single-cell ATAC-seq (scATAC-seq) on 2 representative samples (SML2 and ATL6) to examine the promoter accessibility of *CIITA*. Results showed that the *CIITA* promoter III is more open in infected cells compared to uninfected cells (Supplemental Figure 10D). We then performed a luciferase assay to investigate the effect of different concentrations of Tax on the *CIITA* promoter III activity. We observed that increasing concentrations of Tax caused increasing trans-activation of the *CIITA* promoter III activity (Supplemental Figure 10E). These results suggest that the viral protein Tax plays a role in the upregulation of HLA class II in HTLV-1 infection.

**Figure 5:**
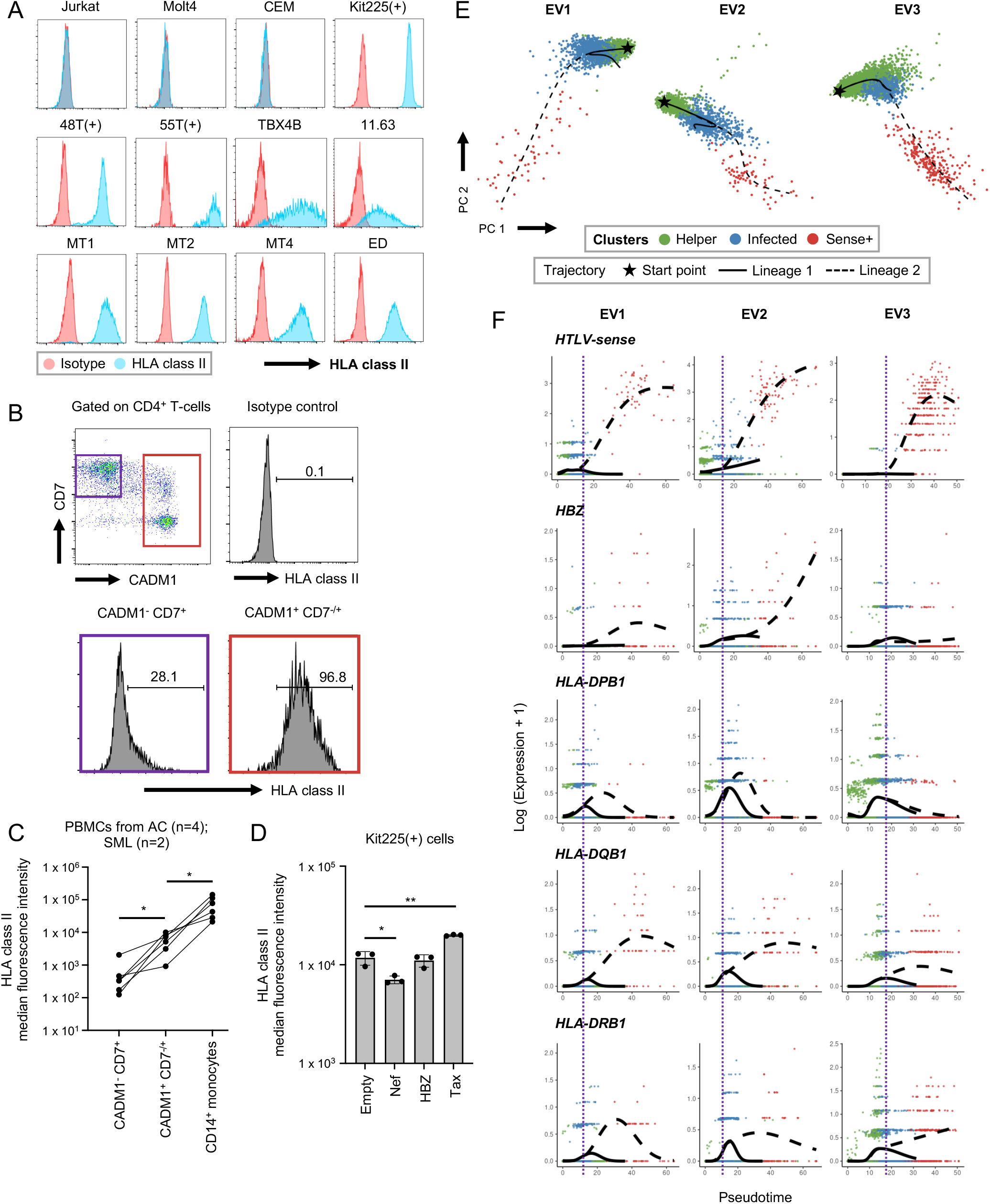
HLA class II expression is associated with HTLV-1-infection and viral protein Tax. (**A**) Plots showing the detection of HLA class II in HTLV-1-infected cell lines. (**B**) Plots show a representative result of the distribution of freshly isolated CD4^+^ T-cells from HTLV-1-infected individuals based on CADM1 and CD7. ATL cells are identified as CADM1^+^ CD7^−/+^ cells and highly express HLA class II molecules. (**C**) Line graph shows the difference of HLA class II expression between non-ATL cells (CADM1^−^ CD7^+^), ATL cells (CADM1^+^ CD7^−/+^) and monocytes (CD14^+^) in 6 HTLV-1-infected individuals. (**D**) Bar graph shows the HLA class II expression in Kit225(+) cells transfected either with an empty vector, HIV-1 nef and HTLV-1 HBZ or tax. (**E**) 2-D PCA plots showing the distribution of helper, infected and HTLV-sense strand expressing cells in 3 different ex vivo cultivated T-cells from HTLV-1-infected individuals. Also shown is the pseudotime trajectories analyzed by Slingshot. (**F**) Expression dynamics of viral and HLA class II genes along the pseudotime trajectories. Color of the dots represent the clusters they are derived from as in panel **E**. The purple line indicates the bifurcation point of the pseudotime trajectories. n = 3 (**D**). Data represent mean ± SD. * p-value < 0.05, ** p-value < 0.01 by unpaired 2-tailed Student’s t-test (**C** and **D**).

### HTLV-1-infected cells can present antigen to responder CD4^+^ T-cells, albeit at a lower efficiency, and induce anergy-related genes

Antigen presentation by professional APCs (B-cells and monocytes) via HLA class II leads to the activation of CD4^+^ T-cells and induces immune response. We aimed to investigate if HTLV-1-infected cells can acquire any ability to present antigens to T-cells when they express HLA class II. It is reported that only certain alleles of the HLA class II genes allow the presentation of endogenous peptides to T-cells, and that HLA-DP molecules with β-chains encoding Gly84 (DP84Gly), such as DP2 and DP4, can present antigen to T-cells(26). Therefore, we used an in vitro model of TCR stimulation through HLA class II utilizing an HLA-DP4-restricted TCR (Clone 9) which is specific to the cancer testis antigen WT1 (Wilms tumor protein-1)(27) to assess the antigen presentation capability of HTLV-1-infected CD4^+^ T cells. In fact, HLA-DP4-expressing and WT1 peptide-pulsed HTLV-1-infected cell lines induced cytokine production in responder T-cells that carry Clone 9 TCR while HTLV-1 negative T-cell lines did not (Figure 6A). Furthermore, peptide-pulsed HTLV-1-infected CD4^+^ T-cells successfully stimulated and induced IFN-γ production in Clone 9-expressing CD4^+^ T-cells, although at a low level, while WT1 peptide-pulsed normal CD4^+^ T-cells and CLIP-pulsed (irrelevant peptide) APCs did not induce any IL-2 production at all (Figure 6, B and C). These results established the ability of HTLV-1-infected CD4^+^ T-cells to present antigen to CD4^+^ T-cells. However, notably, HTLV-1-infected CD4^+^ T-cells as APCs induced a significantly lower number of IFN-γ producing cells in responder T-cells than non-T-cell PBMCs, which are enriched with B-cells and monocytes and commonly used as APCs. This indicates that, although HLA class II-expressing HTLV-1-infected CD4^+^ T-cells can present antigen to T-cells, they are inefficient APCs and not sufficient for fully activating CD4^+^ T-cells (Figure 6C and Supplemental Figure 10, B and C). Next, we aimed to evaluate the effects of antigen-specific stimulation on the expression of key genes for anergy (Figure 6, B and D). We observed that, upon TCR stimulation by HTLV-1-infected CD4^+^ T-cells, the expression of anergy-related molecules was increased compared to when stimulated with non-T-cells containing B-cells and monocytes. (Figure 6D). These results suggest that HTLV-1-infected cells can cause T-cell anergy upon antigen presentation via upregulated HLA class II to escape immunosurveillance in vivo.

**Figure 6:**
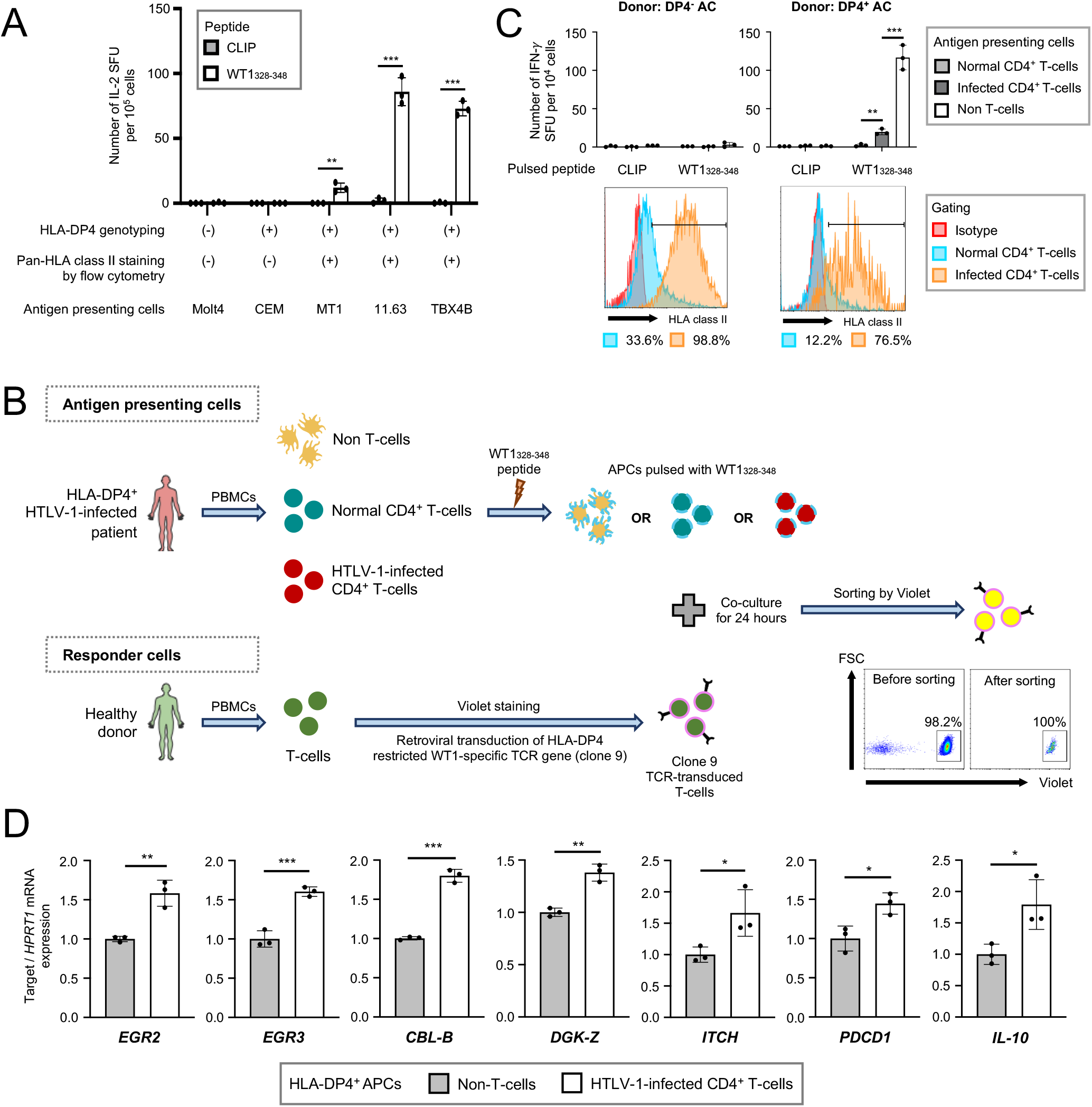
HTLV-1-infected cells can present antigen to responder CD4^+^ T-cells and induce anergy-related genes. (**A**) Bar graph shows the antigen presenting capabilities of different HTLV-1-infected T-cell lines. Jurkat cells expressing HLA-DP4-restricted TCR (clone 9) specific for cancer testis antigen WT1 were stimulated with peptide-pulsed cell lines. CLIP peptide was used as negative control. (**B**) Schematic figure shows the experimental scheme to evaluate T-cell status upon antigen-specific stimulation by HTLV-1-infected CD4^+^ T-cells (**C**) Bar graphs (top row) show the antigen presenting capabilities of different APCs from an HLA-DP4^-^ donor (left) and HLA-DP4^+^ donor (right). Normal CD4^+^ T-cells, infected CD4^+^ T-cells and non-T-cells were prepared from ACs as CADM1^-^ CD4^+^, CADM1^+^ CD4^+^ and CD4/CD8-depleted PBMCs respectively. Clone 9-transduced primary T-cells were stimulated with the APCs pulsed with peptides. Histograms (bottom row) show the expression of HLA class II in normal CD4^+^ T-cells (CADM1^-^ CD4^+^) and infected CD4^+^ T-cells (CADM1^+^ CD4^+^). (**D**) Bar graphs show the expression of T-cell anergy-related genes in responder T-cells after stimulation with non-T-cells (conventional APCs) or HTLV-1-infected CD4^+^ T-cells expressing HLA class II. Non-T-cells and infected CD4^+^ T-cells were prepared from HLA-DP4^+^ ACs as CD3-depleted and CADM1^+^ CD4^+^ PBMCs respectively. Similar results were obtained from at least 2 independent donor samples (**B**). n = 3 (**A** and **C–D**). Data represent mean ± SD. * p-value < 0.05, ** p-value < 0.01, *** p-value < 0.001 by unpaired 2-tailed Student’s t-test (**A** and **C–D**).

## Discussion

Leukemia is a type of cancer originated from blood or bone marrow cells, which is characterized by a large increase in the numbers of abnormal white blood cells. Leukemia can be subdivided into a variety of groups but generally, there are 5 main categories which are acute lymphoblastic leukemia (ALL), acute myeloid leukemia (AML), chronic lymphocytic leukemia (CLL), chronic myeloid leukemia (CML) and other less common types. Inhibition of cell differentiation is a key event of leukemogenesis in ALL and AML. ATL is a leukemia derived from peripheral CD4^+^ T cells. The incidence of peripheral T-cell lymphomas is < 1 case per 100,000 people based on data from the United States Surveillance(28); however, the incidence is around 80 per 100,000 infected individuals in HTLV-1 endemic areas of Japan(29). HTLV-1 infection disrupts the differentiation, activation, proliferation and apoptosis of the host CD4^+^ T-cells, which leads to malignant transformation during its long latency period over decades(10, 13, 30). Thus, it has been extremely difficult to analyze the process of ATL cells generation in the context of CD4^+^ T-cell differentiation/activation dynamics due to its long latency period.

In normal conditions, activated T-cells will induce negative regulatory mechanisms, including FOXP3 and CTLA4, which can terminate T-cell activation and restore homeostasis(31). These homeostatic mechanisms of T-cells are important for preventing excessive inflammation in normal conditions. Our single cell analyses have shown that those key homeostatic mechanisms are actively induced and sustained in HTLV-1-infected cells and ATL cells, which may contribute to immune evasion in the infection. This means that the persistent activation is key for understanding HTLV-1-mediated transformation of T-cells. There are at least three possibilities on how HTLV-1-infected T-cells and ATL cells sustain their highly activated T-cell phenotype with negative regulatory mechanisms operating. Firstly, the viral proteins Tax and HBZ may directly activate mechanisms downstream of TCR signals (e.g. NF-κB complexes) and negative feedback mechanisms for T-cell activation such as transcription factor complexes associated to FOXP3, respectively. Tax enhances NF-κB and AP-1 signaling and contributes to anti-apoptotic and hyper-proliferative phenotype of infected cells(12, 32). On the other hand, HBZ competes with c-Fos for c-Jun binding, inhibiting AP-1 formation downstream of TCR signaling(33). HBZ also induces Treg differentiation by activating TGF-β signaling pathway and thereby inducing FOXP3 expression(34). Secondly, mutations in TCR signaling molecules may sustain activities of genes downstream of TCR signaling as described above(15). Last but not least, HTLV-1 infected T-cells and ATL cells may be enriched with T cells with self-reactive TCRs such as Tregs. It is known that approximately 20% of CD4^+^ T-cells carry self-reactive TCRs and spontaneously receive TCR signals in vivo(18). These three mechanisms are not mutually exclusive and can act synergistically in HTLV-1-infected T-cells, likely operating during leukemic transformation as well. Frequent T-cell over-activation will lead to sustained activation of genes for cell cycle regulation and apoptosis over longer time, which may make infected T-cells more vulnerable to genotoxic stress.

Intriguingly, interferon signaling pathway is actively induced in HTLV-1 infected T-cells and ATL cells by pathway analysis, supporting that anti-viral innate immunity is induced in those T-cells. However, anti-HTLV-1 immunity is actively suppressed in ATL. For example, tax-specific cytotoxic T-cells are reduced in ATL patients compared to HTLV-1 AC(35). Here we propose that the unique dynamics of key surface molecules on ATL cells may contribute to the specific suppression of anti-viral immunity in HTLV-1 infection and anti-leukemia immunity in ATL (immune evasion). Importantly, in our trajectory analysis, T-cells consistently upregulated *CTLA4* throughout the ATL trajectory. CTLA4 efficiently binds to CD80/86 and removes these CD28 ligands from APCs, and thus, inhibits the activities of T-cells and APCs(36). Therefore, the high CTLA4 expression in ATL cells should inhibit T-cell responses in their local microenvironments. In addition, the high expression of HLA class II in ATL cells may contribute to inhibit T-cell responses. T-cells express HLA class II molecules following activation(37, 38), although the functional significance was unknown. Activated Treg (= effector Treg) show enhanced suppressive activities while highly expressing HLA class II molecules(39, 40). Intriguingly, our trajectory analysis showed that HLA class II expression is strongly induced during HTLV-1 infection and further enhanced in ATL. Importantly, our study provides the first demonstration that HTLV-1-infected cells have the ability to present antigen to T-cells. However, HTLV-1-infected cells are not expected to be efficient APCs, because the expression of HLA class II and related molecules in HTLV-1-infected and ATL cells are lower than professional APCs (Figure 3G and Supplemental Figure 10A) and also because HTLV-1-infected cells do not express critical ligands for co-stimulation including CD80/CD86 which conveys the key co-stimulatory CD28 signaling in T-cells (Supplemental Figure 10, B and C). These collectively suggest that HTLV-1-infected cells function and contribute to the escape of infected cells from host immune surveillance (Figure 7), in a similar manner as tolerogenic DCs induce anergy and immunosuppressive function in T-cells(41, 42). Our new model of HTLV-1-infected cells and HLA-DP4-restricted WT1-specific TCR will be useful for future investigations on the molecular mechanism (Figure 6)(26). Summarizing, we have proposed two models on how HTLV-1-infected cells deregulates and evades the host immune system based on the single cell data obtained in this study (Figure 7).

**Figure 7:**
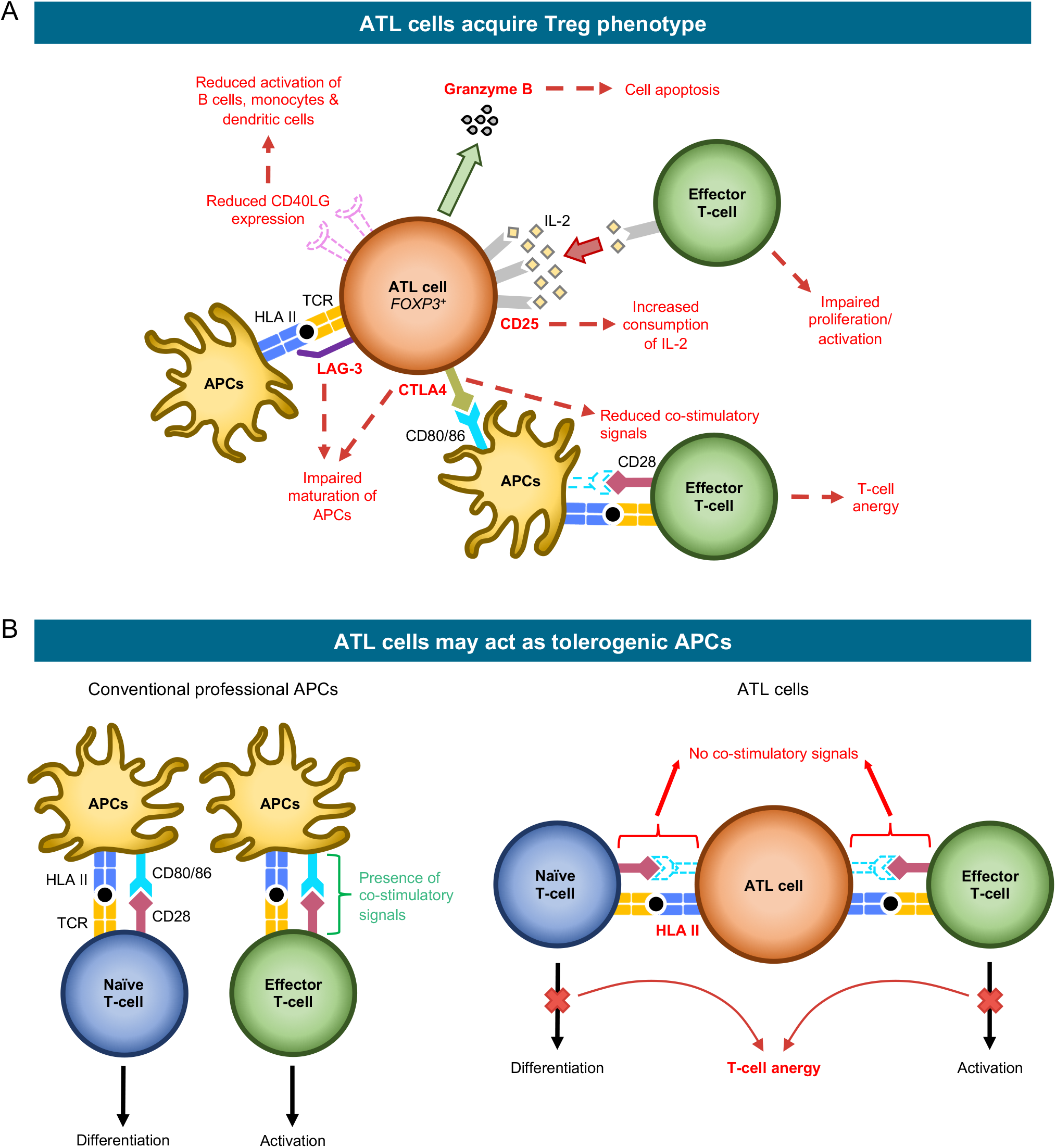
Schematic figure showing our proposed model of the immunosuppressive properties of ATL cells. ATL cells exert their immunosuppressive function by 2 mechanisms: (**A**) Acquiring Treg phenotype. ATL cells upregulate co-inhibitory molecules (*CTLA4, LAG3*) to impair maturation of APCs and reduce co-stimulatory signals required for full T-cell activation. They also upregulate CD25 leading to an increased consumption of IL-2, which reduces the amount of IL-2 required for effector T-cell proliferation and activation; and (**B)** Acting as tolerogenic APCs. ATL cells upregulates HLA class II for antigen presentation but as they lack co-stimulatory signals, target T-cells could not be activated, leading to T-cell unresponsiveness and anergy.

In conclusion, HTLV-1-infection exploits the physiological mechanisms for T-cell activation and homeostasis. Our single cell analysis has provided a new model for HTLV-1 infected CD4^+^ T cells to transform into leukemic cells during the initial phase of ATL leukemogenesis which leads a certain infected clone to malignant transformation in vivo.

## Materials and Methods

Further details of the methods are provided in the supplemental material.

### Patient blood samples

All individuals involved in this study were seen at Imamura General Hospital in Kagoshima city except for the single lymphoma-type ATL patient which was seen at Kumamoto Shinto General Hospital in Kumamoto city. Fresh whole blood was obtained from healthy donors (n = 3) and HTLV-1 infected patients with the following clinical diagnoses: asymptomatic carrier (n = 4), smoldering ATL (n = 3), chronic ATL (n = 6) and lymphoma-type ATL (n = 1). Peripheral blood mononuclear cells (PBMCs) were isolated from whole blood within 24 hours of sample collection using Ficoll-Paque (GE Healthcare) according to the manufacturer’s instructions. Briefly, each blood sample was overlaid on top of Ficoll-Paque at a ratio of 2:1, and centrifuged at 1,800 rpm for 25 minutes at room temperature. Enriched mononuclear cells were washed with PBS and twice centrifuged at 1,600 rpm for 10 minutes. Cell numbers and viability were checked using a hemocytometer and Trypan blue staining.

### Cell suspension preparation

Freshly isolated PBMCs were resuspended in PBS to achieve a cell concentration of 5 × 10^6^ cells/mL. One mL of cell suspension was used for single-cell library preparation and the remaining PBMCs were resuspended in BAMBANKER cryopreservation medium (Nippon Genetics) and stored at -80°C. Prior to single-cell library preparation, dead cells were removed from the 1 mL cell suspension using a dead cell removal kit (Miltenyi Biotec) according to manufacturer’s instructions. After the final wash, cell numbers were determined using a hemocytometer and centrifuged at 1,500 rpm for 5 minutes. The resulting cell pellet was then resuspended in PBS plus 0.04% BSA to a concentration of 1,000 cells/µL.

### Proviral load measurement of clinical samples by droplet digital PCR

Genomic DNA was extracted from 5 × 10^6^ cells using Qiagen DNeasy Blood & Tissue Kit (Qiagen) following manufacturer’s instructions. Fifty or 100 ng genomic DNA were used for each droplet digital PCR (ddPCR) reaction. ddPCR was performed using primers and a probe targeting a conserved region in HTLV-1 pX region and the *ALB* gene as was previously reported(43). Droplet quantification was performed on a QX200 droplet reader (Bio-Rad) and data analyzed using QuantaSoft software (v1.7.4, Bio-Rad). For an objective cut-off with maximum sensitivity, a no-template-control (NTC) sample was used to determine the threshold values in which the highest level of droplet fluorescence in NTC was designated as the threshold line for negative signal. Proviral load (PVL) was thus calculated as follows: PVL(%) = 100 × (HTLV copy number × 2)/ALB copy number.

### Single-cell library preparation

Live cells-enriched cell suspensions were loaded onto a Chromium Single Cell Chip (10× Genomics) for co-encapsulation with barcoded gel beads from the Single Cell 5’ Gel Bead Kit (10× Genomics) at a target capture rate of 10,000 individual cells per sample. Barcoded cDNAs were pooled, PCR amplified and then used to generate 2 different libraries. A single-cell transcriptome library was prepared with 50 ng of cDNA amplified product using the Chromium Single Cell 5’ Library Kit (10× Genomics). Enrichment of V(D)J segments and construction of single-cell TCR library was performed using the Chromium Single Cell V(D)J Enrichment Kit, Human T Cell and the Chromium Single Cell 5’ Library Construction Kit respectively (10× Genomics). The libraries were sequenced with paired-end, single indexing at 150 + 150 bp reads (TCR library) or 26 + 91 bp reads (transcriptome library) on the Illumina HiSeq or NextSeq platform.

### Canonical correspondence analysis (CCA)

Explanatory variables for CCA were prepared as follows. The T-cell activation explanatory variable was defined by the differentially expressed genes (DEGs), obtained using the Bioconductor package DESeq2 (v1.30.1), between TCR-stimulated only T-cells (Th0) and resting T-cells from a published scRNA-seq dataset of CD4^+^ T-cells (44) (https://www.opentargets.org/projects/effectorness). The regulatory T-cell (Treg) explanatory variable was defined by the DEGs, obtained using GEO2R (https://www.ncbi.nlm.nih.gov/geo/geo2r/), between resting Treg and resting conventional T-cells (Tconv) from GSE15390(45). For both explanatory variables, genes with FDR < 0.05 and log2 fold change (> 0.5 or < -0.5) were selected. For the 1D CCA, the expression data of CD4^+^ T-cells only were regressed onto each of the explanatory variables above and correspondence analysis was performed using the Bioconductor package vegan (v2.5-7). For the 2D CCA, the expression data of CD4^+^ T-cells only were regressed onto both explanatory variables followed by correspondence analysis. Regression model of the CCA scores were obtained by applying a generalized additive model to the dataset using the Bioconductor package ggplot2 (v3.3.5).

### Pseudotime analysis

Pseudotime trajectories were identified using the monocle package (v2.18.0) (46). We prepared the Seurat object for pseudotime analysis by first excluding all cytotoxic T-cells (C1-C3) from the Seurat object. Next, we removed cells which expressed CD8A/CD8B and also cells with unidentified clonotype in TCR-seq. For the analysis involving the entire T-cell population, we converted the resulting Seurat object into a CellDataSet (CDS) object and pass it to monocle for pseudotime analysis. For the individual / paired ATL analysis, we performed another sub setting where we kept only cells from the three healthy individuals and the ATL patient of interest before converting it to a CDS object. Clusters which are enriched with CCR7^+^ cells are assumed to be the origin of the trajectory. Genes which varied as a function of pseudotime were identified using monocle’s “differentialGeneTest” function.

### RNA velocity analysis

RNA velocity analysis was performed on the dataset of 3 HDs and paired ATL sample. Using the aligned BAM files from Cell Ranger, the number of spliced and unspliced reads were recounted using the Python package velocyto (v0.17.17)(47) to generate the loom files. To generate the UMAP and clusters for these dataset, we first subset these 5 datasets from the Seurat object of T-cells. Next, the top 3,000 variable features were detected followed by removal of mitochondrial, ribosomal and T-cell receptor genes. Data integration was performed with Harmony and UMAP was used for visualization of unsupervised clustering. Dynamical model velocities were computed from the loom files and visualized on UMAP using the Python package scVelo (v0.2.3)(48).

### TCR-seq analysis

TCR sequences were assembled with the Cell Ranger vdj pipeline (v3.1.0, 10× Genomics), leading to an output file which contains the identification of CDR3 sequence and rearranged TCR gene. Cells with identified clonotype (i.e. not NA or ‘none’) are kept while the others were discarded. Clonality was then designated as expanded for cells with 2 or more similar TCR while the remaining ones are marked as unexpanded.

### Quantification of HTLV-1 proviral load in sorted cells

Measurement of HTLV-1 proviral load (PVL) of sorted cells (CADM1^-^ CD7^+^ and CADM1^+^ CD7^-/+^ cells) was as described previously(21). Briefly, quantitative multiplex real-time PCR was performed with two sets of primers specific for the HTLV-1 provirus and the human gene encoding the RNase P enzyme. PVL was expressed as HTLV-1 copy numbers per 100 PBMCs based on the assumption that infected cells harbor one copy of integrated HTLV-1 provirus per cell.

### Targeted exome sequencing and analysis

Target capture was performed using a HTLV-1/ATL panel and the SureSelect Target Enrichment System (Agilent Technologies) as reported previously(49). Resulting libraries were sequenced with 100 bp paired-end reads on the Illumina HiSeq platform. Sequenced data were aligned to the human reference genome hg38 using BWA (v0.7.15) software. PCR duplicates were removed using Picard (v2.92) and SAMtools (v1.2) softwares. Uninfected T-cells (CD4^+^CADM1^-^CD7^+^) were used as matched normal controls to call somatic mutations. Somatic mutation candidates were called using MuTect2 from the GATK (v4.0.12) software and annotated with ANNOVAR (v20191024). Candidate mutations with (i) ≥5 variant reads in tumor samples, (ii) a VAF in tumor samples ≥ 0.01, (iii) read depth ≥200, and (iv) tumor:normal variant ratio ≥2 were adopted and further filtered by excluding synonymous SNVs.

### Preparation of PBMCs and cell lines for flow cytometry analysis

PBMCs were obtained via density gradient centrifugation using Ficoll-Paque PLUS (GE Healthcare Life Sciences). The leukemic T-cell lines Jurkat, Molt4 and Kit225(+) were obtained from ATCC (TIB-152), ATCC (CRL-1582) and ATCC (CRL-1990) respectively. CEM, a leukemic T-cell line, was obtained from Dr. Hiroaki Takeuchi. The ATL cell lines ATL-48T(+), ATL-55T(+), MT1 and ED were obtained from Dr. Michiyuki Maeda (Maeda M. et al., J Exp Med. 1985). The HTLV-1-infected cell lines MT-2, MT-4, TBX4B, and 11.63 were obtained from Prof. Charles R.M. Bangham(50). IL-2 dependent cell lines were cultured in the presence of recombinant IL-2 (100 U/ml). All T-cell lines were cultured in RPMI 1640 supplemented with 10% fetal bovine serum (FBS) and penicillin/streptomycin.

### Analysis of ex vivo cultivated primary cells

Fresh PBMCs were cultivated in RPMI 1640 supplemented with 10% FBS and penicillin/streptomycin for 18 hours to induce viral sense strand transcription in naturally infected cells. Harvested cells were subjected to single cell library preparation and resulting scRNA-seq data were analyzed as mentioned previously. Pseudotime trajectories were identified using the Bioconductor package slingshot (v1.8.0)(51) and projected onto 2-D PC space. Clusters which are enriched with CCR7^+^ cells are assumed to be the origin. A negative binomial generalized additive model (GAM) for each gene is fitted using the Bioconductor package tradeSeq (v1.4.0)(52).

### Flow Cytometry Analysis

The following reagents were used for flow cytometry analysis: BV510-conjugated anti-HLA-II (Tu39, BioLegend), BV421-conjugated anti-CD7 (M-T701, BD Biosciences), anti-CADM1 (3E1, MBL), FITC-conjugated anti-CD80 (2D10, Biolegend), PE-conjugated anti-CD86 (IT2.2, Biolegend), polyclonal Alexa647-conjugated goat anti-chicken IgY (Abcam), PE-conjugated Streptavidin (BioLegend), PerCP/Cyanine5.5-conjugated anti-CD14 (HCD14, BioLegend), PerCP/Cyanine5.5- or biotin-conjugated anti-CD4 (OKT4, BioLegend). Dead cells were distinguished with LIVE/DEAD Fixable Near-IR Cell Stain Kit (Thermo Fisher Scientific). Primary ATL cells and cell lines were pre-treated with goat serum for 20 minutes at room temperature. After washing, cell surface molecules were stained with specific antibodies for 20 minutes at 4 °C. Stained cells were analyzed with FACSVerse (BD Biosciences). Data analysis was performed using FlowJo software (version 10.7.1) (Tree Star).

### Transfection of *tax* and *HBZ*

HTLV-1 viral gene expression vectors such as *tax* and *HBZ* were generated based on pIRES-EGFP (6029-1, Clontech). As a negative control, HIV-1 n*ef* gene expression vectors was also generated based on pIRES-EGFP. Kit225(+) cells were transfected with each viral expression vector by electroporation using NEPA21 (NEPAGENE).

### Assessment of antigen presentation by upregulated HLA class II in HTLV-1-infected cells

For the ELISPOT assay, all HTLV-1-infected cell lines and donors were typed for HLA-DP at the HLA Foundation Laboratory. TCR-deficient Jurkat cell line was established previously(53). Synthetic peptides (CLIP: LPKPPKPVSKMRMATPLLMQALPM and WT1_328-348_: PGCNKRYFKLSHLQMHSRKHT) were purchased from Genscript and dissolved at 50 mg/ml in DMSO. Primary T-cells were purified using the Pan T-cell Isolation Kit (Miltenyi Biotec). HLA-DP4 restricted WT1 specific TCR (clone 9)(27) were transduced into TCR-deficient Jurkat cells or primary T-cells using the Plate-GP and PG13 cell-based retrovirus system and Retronectin (Takara Bio) according to manufacturer’s instructions. HLA-DP4/WT1 TCR-transduced Jurkat cells were stimulated with HTLV-1-infected cell lines pulsed with WT1_328-348_ (10 µg/ml) at an E:T ratio of 5:1 for 24 hours followed by measurement of specific cytokine production using the human IL-2 ELISpot kit (Mabtech AB) and ELIPHOTO Counter (Minerva Tech).

For stimulation of TCR-transduced primary T-cells, CD4^+^ T-cells were first isolated from HLA-DP4^+^ asymptomatic carriers using the Pan T-cell Isolation Kit (Miltenyi Biotec) and biotin-labeled anti-CD8 mAb (RPA-T8, BioLegend). Next, HTLV-1-infected cells were sorted as CD4^+^CADM1^+^ T-cells using FITC-labeled anti-CADM1 mAb (3E1, MBL) and anti-FITC MicroBeads (Miltenyi Biotec). The CADM1^-^ fractions were used as normal CD4^+^ T-cells. Non-T-cells containing monocytes and B-cells were prepared by T-cell depletion using anti-biotin MicroBeads (Miltenyi Biotec), biotin-labeled anti-CD4 mAb (OKT4, BioLegend) and biotin-labeled anti-CD8 mAb (RPA-T8, BioLegend). Primary T-cells (1 × 10^4^ cells) expressing clone 9 were stimulated for 24 hours with HLA-DP4^+^ non-T-cells, normal CD4^+^ T-cells or HTLV-1-infected CD4^+^ T-cells (2 × 10^4^ cells) pulsed with WT1_328-348_ (10 µg/ml) followed by measurement of specific cytokine production using the human IFN-γ ELISpot kit (Mabtech AB) and ELIPHOTO Counter (Minerva Tech).

### Assessment of T-cell anergy status upon antigen-specific stimulation by HTLV-1-infected cells

Responder T-cells were generated by transducing HLA-DP4/WT1 TCR (clone 9) into primary T-cells as shown in Figure 6C and labeled with a tracer dye (CellTrace Violet, Life Technologies). HTLV-1-infected CD4^+^ T-cells from HLA-DP4^+^ asymptomatic carriers were as purified as shown in Figure 6C. Non-T-cells were prepared by T-cell depletion using anti-FITC MicroBeads (Miltenyi Biotec) and FITC-labeled anti-CD3 mAb (SK7, BioLegend). Both stimulators were pulsed with WT1_328-348_ (10 µg/ml) for 2 hours and then co-cultured with responder T-cells for 24 hours. Cells were then stained with LIVE/DEAD Fixable Near-IR Cell Stain Kit (Thermo Fisher Scientific) and Violet^+^ cells (responder T-cells) were sorted out using SH800S Cell Sorter (SONY). Quantification of genes associated with T-cell anergy were performed by real-time PCR. cDNA was synthesized using ReverTra Ace qPCR RT Master Mix (Takara Bio) followed by quantification using the THUNDERBIRD SYBR qPCR Mix (TOYOBO) and StepOnePlus Real-Time PCR System (Applied Biosystems). Primer sequences for the genes used are listed in Supplementary Table 2. The relative expression level for each sample was calculated using the ΔΔCt method with the expression level of *HPRT1* in CLIP peptide-stimulated T-cells as the reference sample.

### Statistics

Statistical analyses for CCA scores were performed with R (v4.0.3) using Wilcoxon rank sum test and p-values adjusted using Bonferroni-Hochberg correction for multiple testing. Statistical analyses for flow cytometric, ELISPOT and real-time PCR data were performed using GraphPad Prism 7 software (GraphPad Software). Unpaired 2-tailed Student’s t-test were used for two sample comparisons. A p-value of less than 0.05 was considered statistically significant.

### Study approval

Study participants gave written informed consent prior to blood collection. The study was approved by the ethical review committee of Kumamoto University (Genome No. 263).

## Supporting information

Supplemental infomation

## Data availability

The expression matrix and associated metadata can be assessed in GEO under the accession number XXX. The source code to reproduce our analyses can be accessed at XXX.

## Author contributions

BJYT, MO and Y Satou conceptualized and designed the study; BJYT, KS, OR, MM, K Uchiyama, PM, MY, Y Suzuki and HK performed the experiments; BJYT, KS, VH, MY, K Uchimaru, YU, MO and Y Satou analyzed the data; TU provided materials; HS, MT, HN, ES and AU took care of patients, procured blood samples and provided clinical information; BJYT, MO and Y Satou wrote and edited the manuscript. All authors read and approved the final manuscript.

## Acknowledgments

We thank all the donors who provided blood samples for this study. We also thank Dr. David Robertson (University of Glasgow) for valuable discussion about bioinformatics analysis. This work was supported by research grants from the Japan Society for the Promotion of Science (JSPS) KAKENHI (JP20H03724 and JP18KK0230 to Y Satou; JP19H05426, JP21K07082, and JP21H00433 to MO; 16KK0206 and JP18K16122 to HK; JP18K08437 and JP18KK0452 to PM; JP20K22783 and JP21K08494 to KS; JP21K15454 to MM); Japan Agency for Medical Research and Development (AMED) (JP20jm0210074, JP20wm0325015, JP19fm0208012 and JP20fk0410023 to Y Satou); the Grant for Joint Research Project of the Institute of Medical Science, the University of Tokyo to Y Satou; the grant from Kumamoto University Excellent Research Projects to Y Satou and JST MIRAI (18077147) to Y Satou; the program of the Joint Usage/Research Center for Developmental Medicine, Inter-University Research Network for Trans-Omics Medicine, Institute of Molecular Embryology and Genetics, Kumamoto University to Y Satou. BJYT is a PhD candidate at Kumamoto University and this work is submitted in partial fulfilment of the requirement for the PhD.

