## Supplemental infomation for "Retrovirus-induced leukemia – hijack of T-cell activation mechanisms revealed by single-cell analysis"

### Supplemental Methods

#### scRNA-seq data processing and analysis

Sequencing data alignment and gene quantification were performed with the Cell Ranger Single-Cell Software Suite (v3.1.0, 10x Genomics) using a modified hg38 human reference genome which contains the HTLV-1 genome (Genbank accession no. AB513134) as a separate chromosome. The resulting output file, which is a gene-cell barcode matrix, was imported into R (v4.0.3) and analyzed using Seurat (v4.0.3) (1). Before proceeding with downstream analysis, ambient mRNA contamination was estimated using hemoglobin genes and removed from the data using SoupX (v1.5.2) (2) while doublets were predicted *in silico* and removed using scDblFinder (v1.4.0) (https://github.com/plger/scDblFinder). This is next followed by a 2-step filtering to remove low quality cells. First, cells with less than 200 genes detected or any genes which are expressed in less than 3 cells were removed from the data set. Next, cell quality was evaluated based on three metrics: (1) The total number of UMI counts per cell; (2) The number of genes detected per cell; and (3) The proportion of mitochondrial genes detected. Cells fulfilling the following criteria are deemed to be of high-quality and retained for downstream analysis: (1) The total number of UMI counts per cell or proportion of mitochondrial genes detected was below the mean of all cells plus 2 × standard deviation; and (2) The number of genes detected was within the mean of all cells ± 2 × standard deviation. Counts were then normalized and log-transformed using Seurat's default normalization parameters. The top 3,000 variable genes were detected using Seurat's

“FindVariableFeatures” function followed by removal of mitochondrial and ribosomal genes from the list. Principal component analysis (PCA) was performed on the scaled data followed by batch correction with Harmony (v0.1.0) (3). A uniform manifold approximation and projection (UMAP) 2-dimensional representation of the data was calculated using the first 30 harmonized dimensions for visualization. Clusters were calculated using Seurat’s “FindClusters” function with default parameters. Cluster annotation was performed based on a cell module score calculated using Seurat’s “AddModuleScore” function.

**T-cell re-integration and analysis**

For T-cell analysis, we first subset the T-cell clusters from the Seurat object. To ensure that only pure T-cells were retained, we retained only cells having a T-cell module score which is higher than the maximum score of non-CD3 (CD3D, CD3E and CD3G) expressing cell in the dataset. Following that, we detected the top 3,000 variable features and removed mitochondrial, ribosomal and T-cell receptor genes. Data integration was performed with Harmony and UMAP was used for visualization of unsupervised clustering. Module scores for the activation and exhaustion signature was calculated using Seurat’s “AddModuleScore” function. Clusters were annotated based on examination of T-cell-related genes, HTLV-1-related genes (*HTLV-sense*, *HBZ*, *CADM1*, *CCR4*, *CD40LG*, *CD7* and *DPP4*) and T-cell clonality. Reference mapping was also performed to check our annotations. This was done by mapping our dataset using default parameters to an annotated dataset of 162,000 PBMCs in Seurat(4).

**Differential gene expression and pathway analysis**

Differential expression testing between different clusters and groups were performed with Seurat’s “FindMarkers” function using the MAST (Mode-based Analysis of Single-cell Transcriptomics) method(5). For each HTLV-1-infected / ATL cluster, DEGs were generated relative to cluster H2. Pathway analysis was conducted with the Bioconductor package clusterProfiler (v3.18.1) (6) using the Reactome database.

**scATAC-seq library preparation, data processing and analysis**

Nuclei were isolated from cryopreserved PBMCs ($5 \times 10^6$) using the recommended protocol by 10x Genomics (CG000169: Nuclei Isolation for Single Cell ATAC Sequencing RevD). Nuclei pellet was then resuspended in Diluted Nuclei Buffer to a concentration of 5,000 nuclei/μL for a targeted nuclei recovery of 10,000 nuclei. Transposition, single-cell droplet generation and library construction were then performed using the Chromium Next GEM Single ATAC Library & Gel Bead Kit (10x Genomics) according to manufacturer's protocol. Libraries were sequenced with paired-end, dual indexing at 50 + 50 bp reads on the Illumina HiSeq platform. Sequencing data alignment and peak calling were performed with the Cell Ranger ATAC (v1.2.0, 10x Genomics) using the hg19 human reference genome. The resulting output file, which is a peak-cell barcode matrix, was imported into R (v4.0.3) and analyzed using Signac (v1.3.0)(7). Standard analysis methods were performed in accordance to the vignette on Signac's webpage available here (https://satijalab.org/signac/articles/pbmc_vignette.html). To facilitate interpretation of scATAC-seq data, the corresponding scRNA-seq data was integrated with the scATAC-seq data which enables us to identify the clusters in scATAC-seq based on the annotations in the scRNA-seq data. A peak coverage plot was then generated to evaluate chromatin accessibility across the CIITA region (chr16:10,960,000–11,035,000).

**Luciferase assay**

Nucleotides -322 to +101 of the human CIITA promoter-III were cloned into pGL4.10 (Promega)(8, 9). Jurkat cells ($2 \times 10^5$) were transfected with a reporter plasmid, a Renilla luciferase control vector (pRL-TK) and a Tax-expression plasmid (pCG-Tax) using Lipofectamine LTX with Plus Reagent (Thermo Fisher Scientific) and harvested after 24 hours. Luciferase assays were then performed using the Dual-Glo Luciferase Assay System (Promega) according to manufacturer's instructions and luminescence was detected using GloMax Luminometer (Promega). Relative luciferase activities were calculated as the ratio of firefly to Renilla luciferase activities.

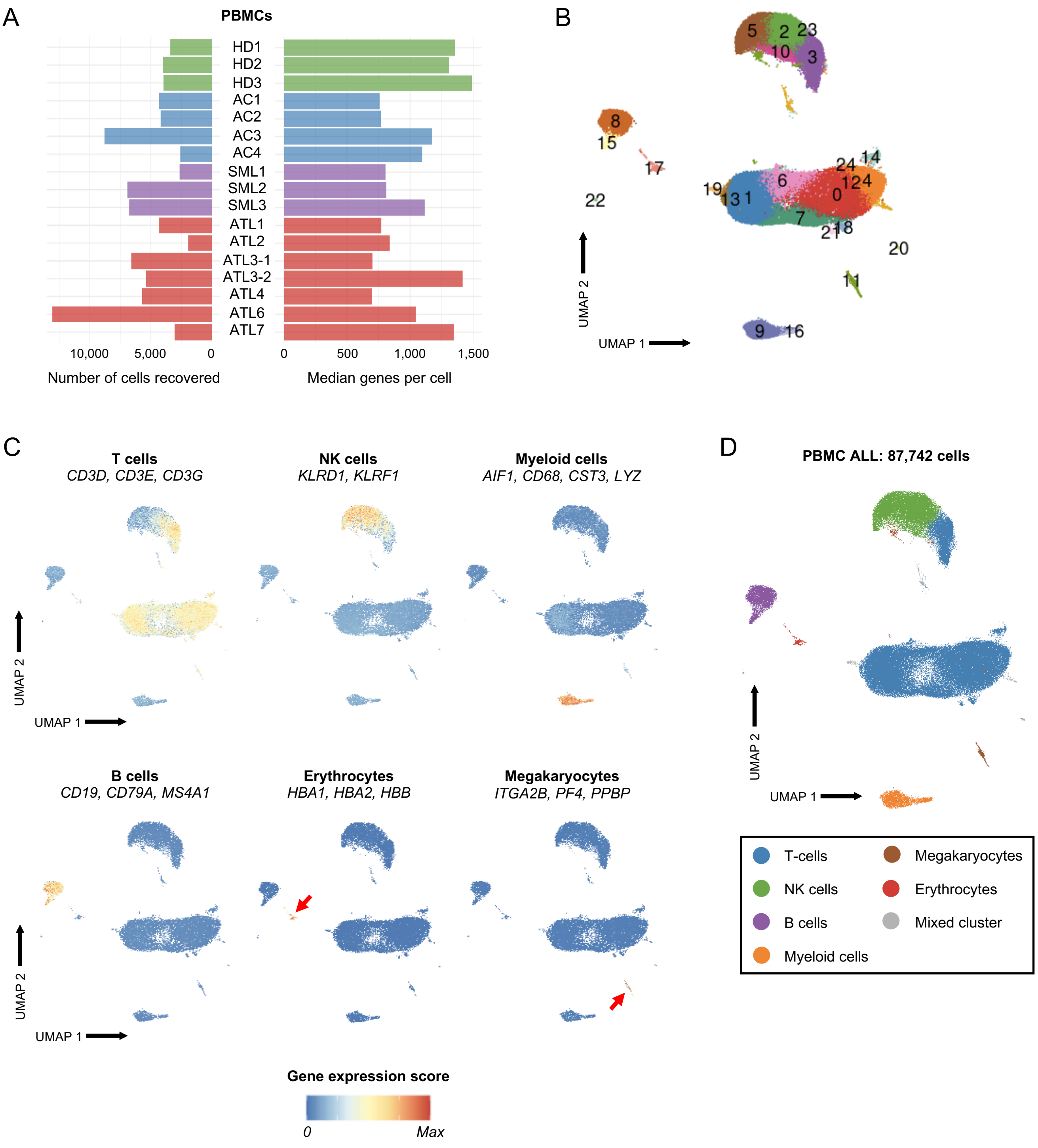


**Figure S1: Identification of PBMC sub-populations.** (**A**) The number of PBMCs recovered that passed quality controls (left) and their median number of genes per cell (right) for each of the individuals (3 healthy donors, HD1-HD3; 4 asymptomatic carriers, AC1-AC4; 3 smoldering ATL, SML1-SML3; and 7 ATL, ATL1-ATL7). (**B**) 2-D UMAP plot shows the 25 clusters identified by Seurat for the entire PBMC population. (**C**) 2-D UMAP plots show the gene module score for each PBMC sub-population. (**D**) Plot shows the 7 sub-populations that can be identified in the 2-D UMAP of entire PBMC population as in panel **B**.

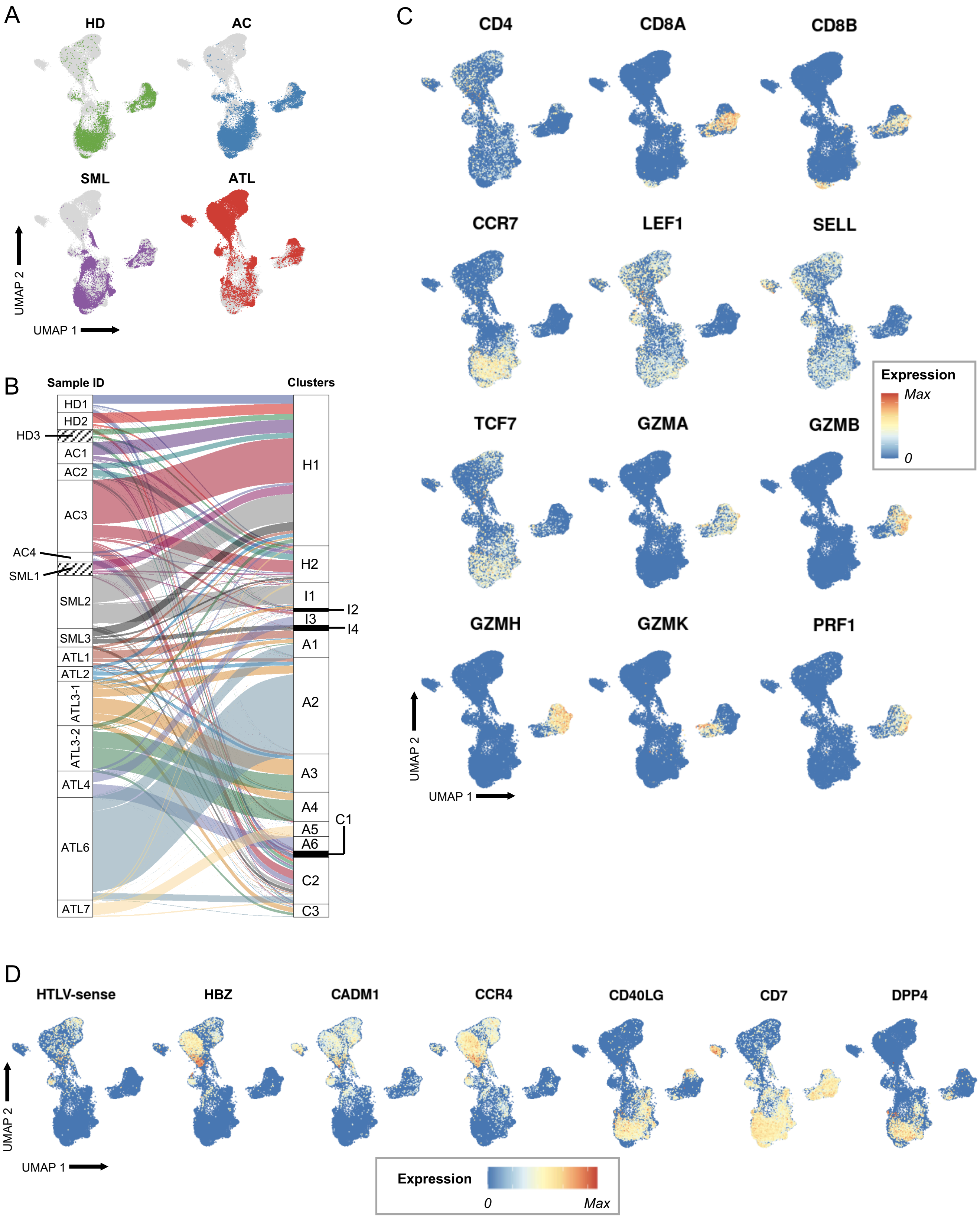


**Figure S2: Identification of T-cell sub-populations.** (**A**) Plots show the distribution of each clinical group in the 2-D UMAP plot in Figure **1C**. (**B**) Alluvial plot shows the assignment of each sample to each cluster as identified in Figure **1C**. (**C**) Plots show the expression of T-cell-related genes. (**D**) Plots show the expression of genes related to HTLV-1 infection.

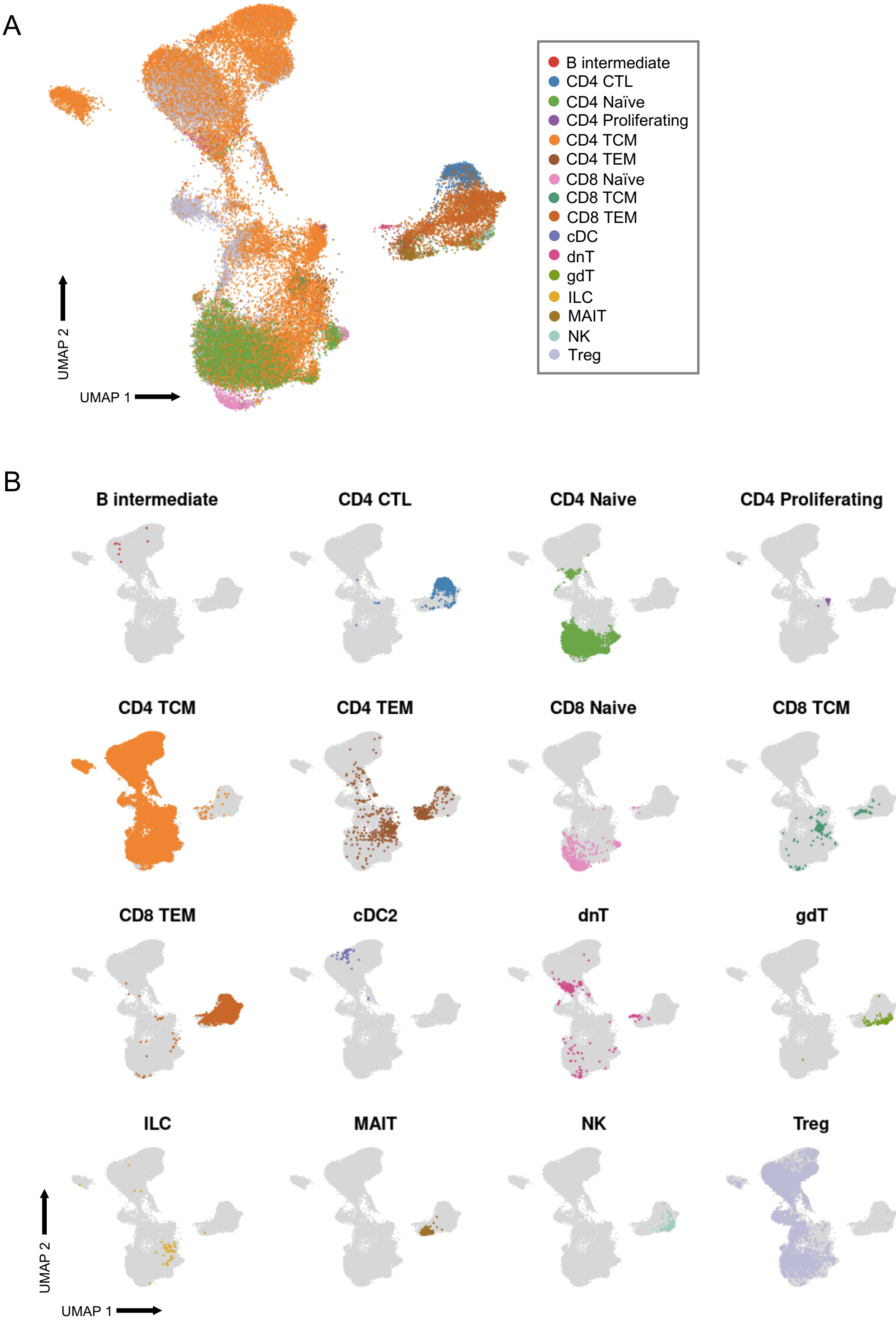


**Figure S3: T-cell sub-populations identified by reference mapping.** (**A**) 2-D UMAP plot shows the single cell annotations by mapping to a reference dataset. (**B**) Plots show the distribution of the cell identities shown in panel **A** in the 2-D UMAP plot.

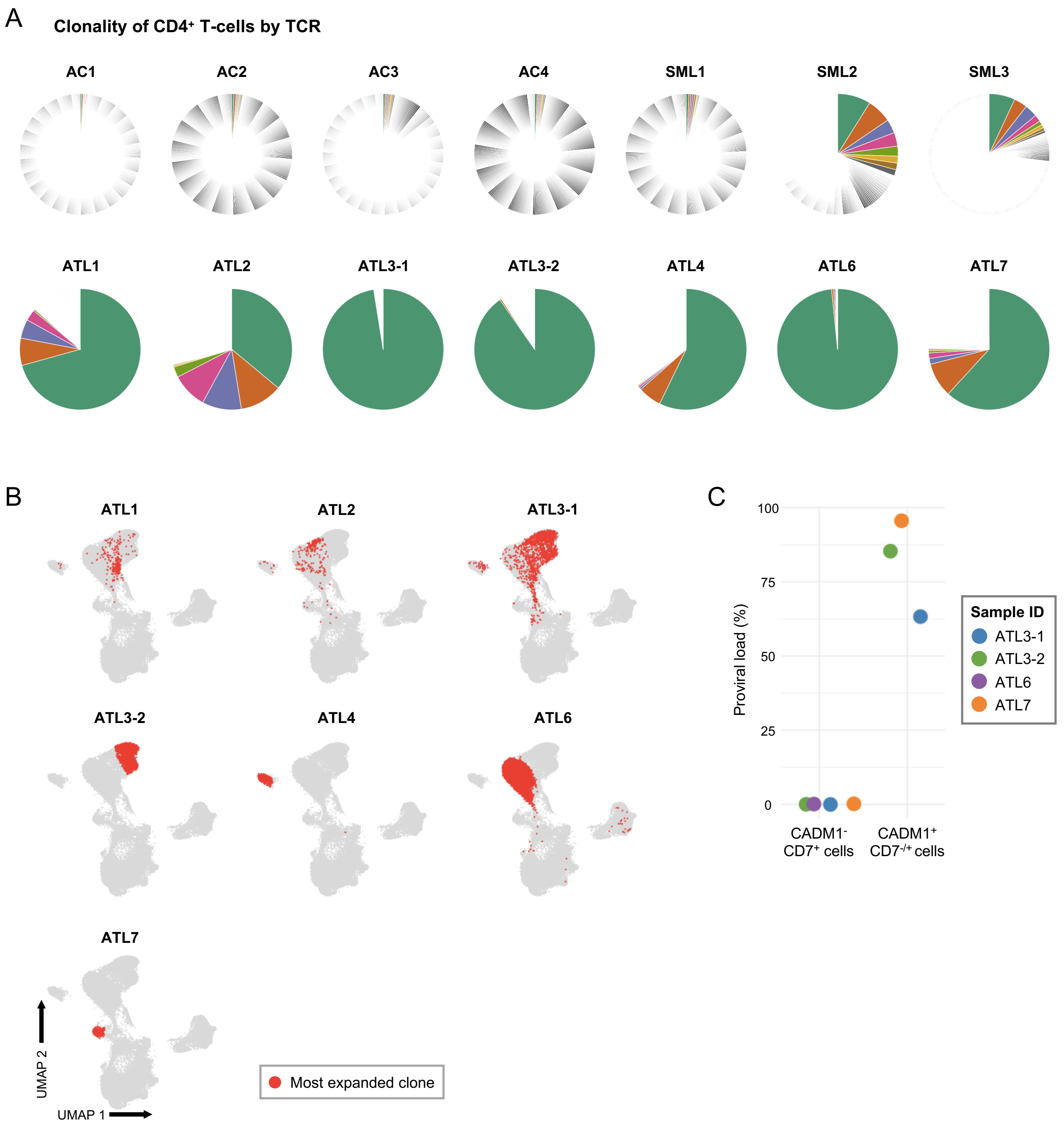


**Figure S4: CD4⁺ T-cells clonality and proviral load.** (**A**) Pie charts show the distribution of T-cell clonality identified through TCR-seq in CD4⁺ T-cells. (**B**) Plots show the distribution of the most expanded clone for each ATL patient in 2-D UMAP space. (**C**) Dot plot shows the proviral load of ATL cells (CADM1⁺ CD7⁻/⁺) and non-ATL cells (CADM1⁻ CD7⁺).

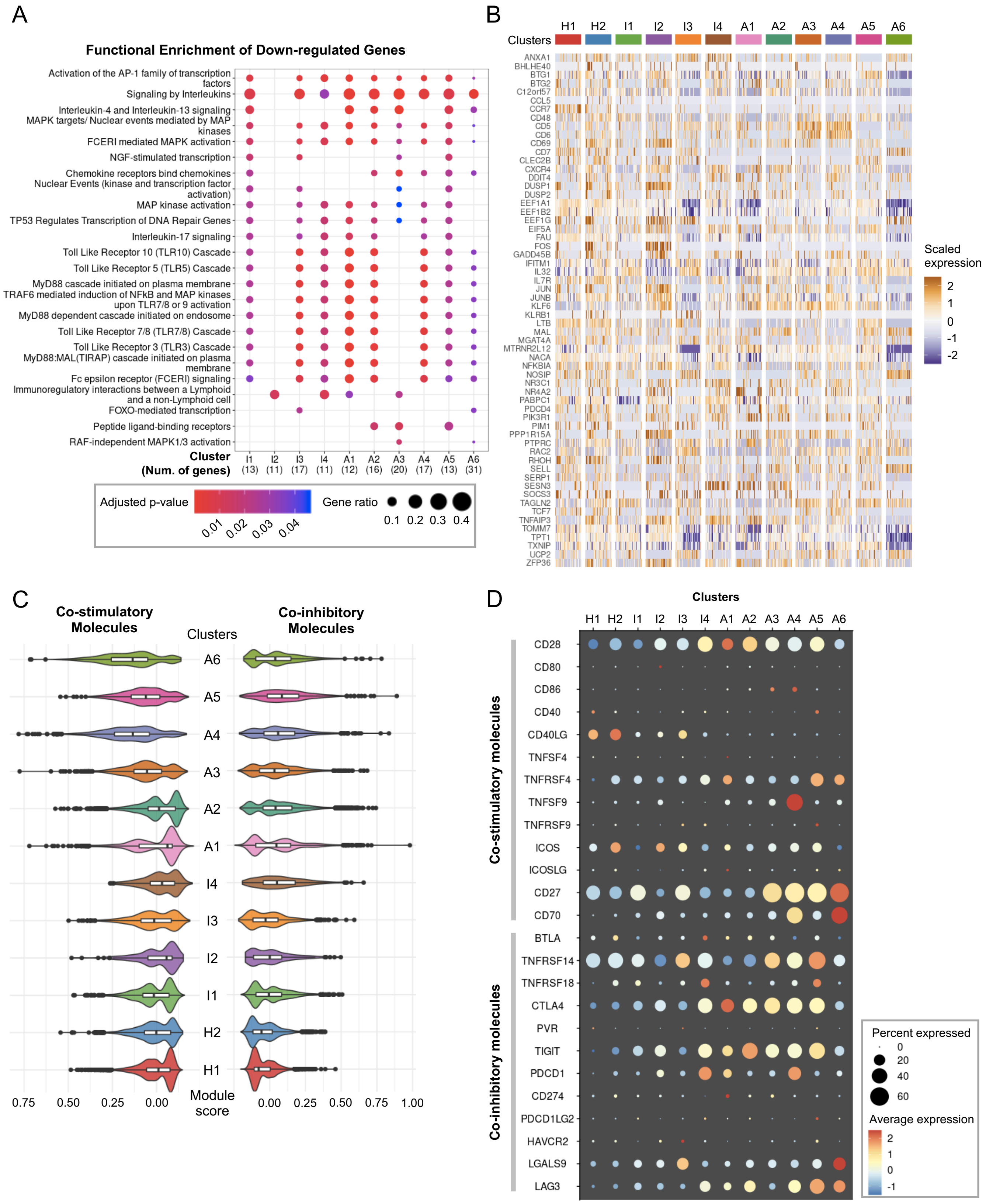


**Figure S5: ATL cells highly express co-stimulatory receptors but showed no common downregulated genes.** (**A**) Dot plot shows the shared and distinct Reactome pathways of downregulated genes for each infected cluster. (**B**) Heatmap showing the expression of downregulated genes in each cluster. (**C**) Violin plot shows the distribution of co-stimulatory (left) and co-inhibitory (right) gene module scores for all CD4$^+$ T-cell clusters. Box plots in each violin summarize the median (midline) and IQRs. (**D**) Dot plot shows the average expression and percentage of expressing cells of each gene in the activation and exhaustion signature for all CD4$^+$ T-cell clusters.

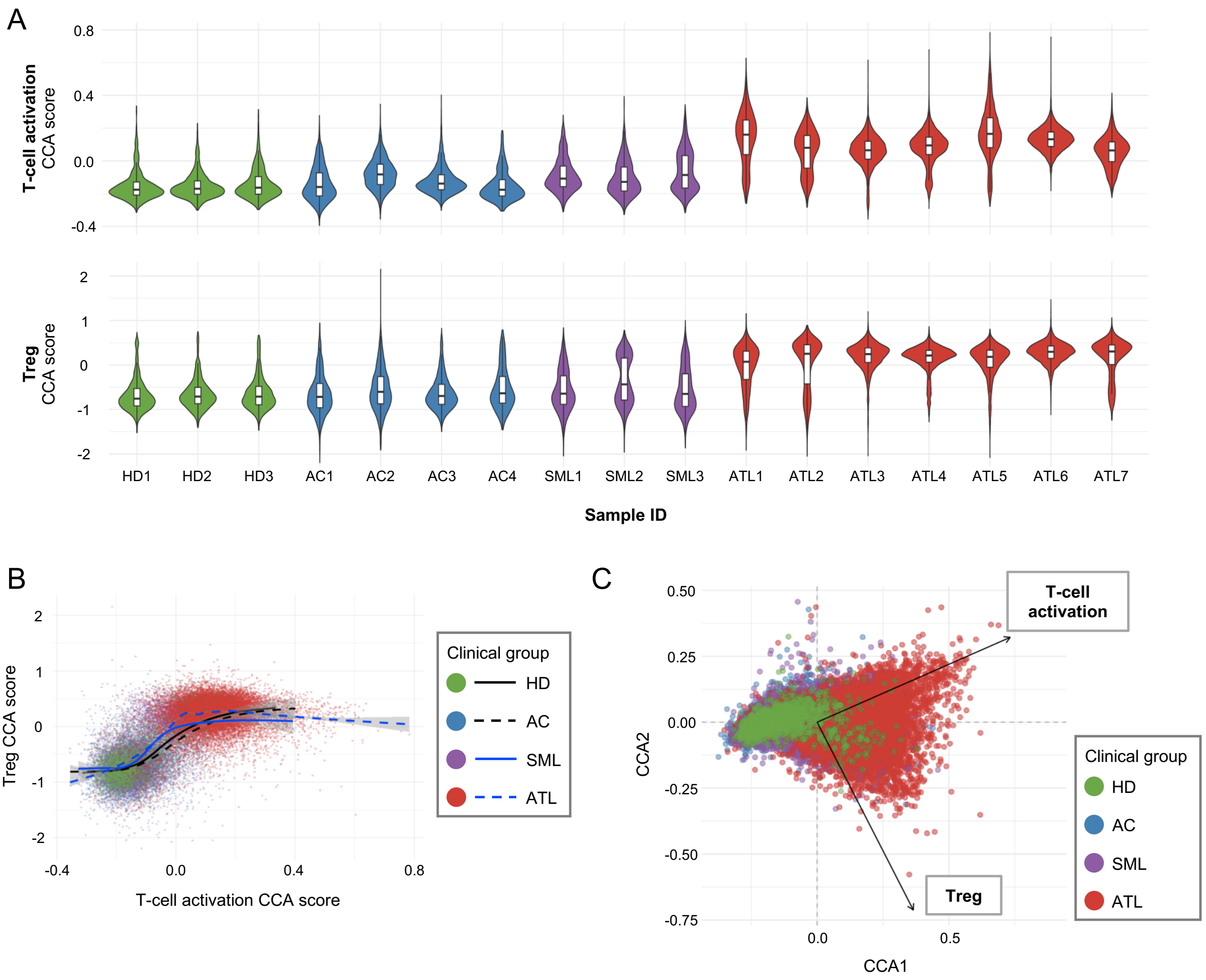


**Figure S6: CCA analysis.** (**A**) Violin plots show the 1-D CCA scores for T-cell activation and Treg grouped by sample ID. Box plots in each violin summarize the median (midline) and IQRs. (**B**) Scatter plot shows the correlation between the 1-D CCA score for T-cell activation and Treg. Lines show the regression model of the CCA scores for all different clinical groups. (**C**) Scatter plot shows the 2-D CCA of all CD4$^{+}$ T-cells. CCA1 is for T-cell activation and CCA2 is for Treg.

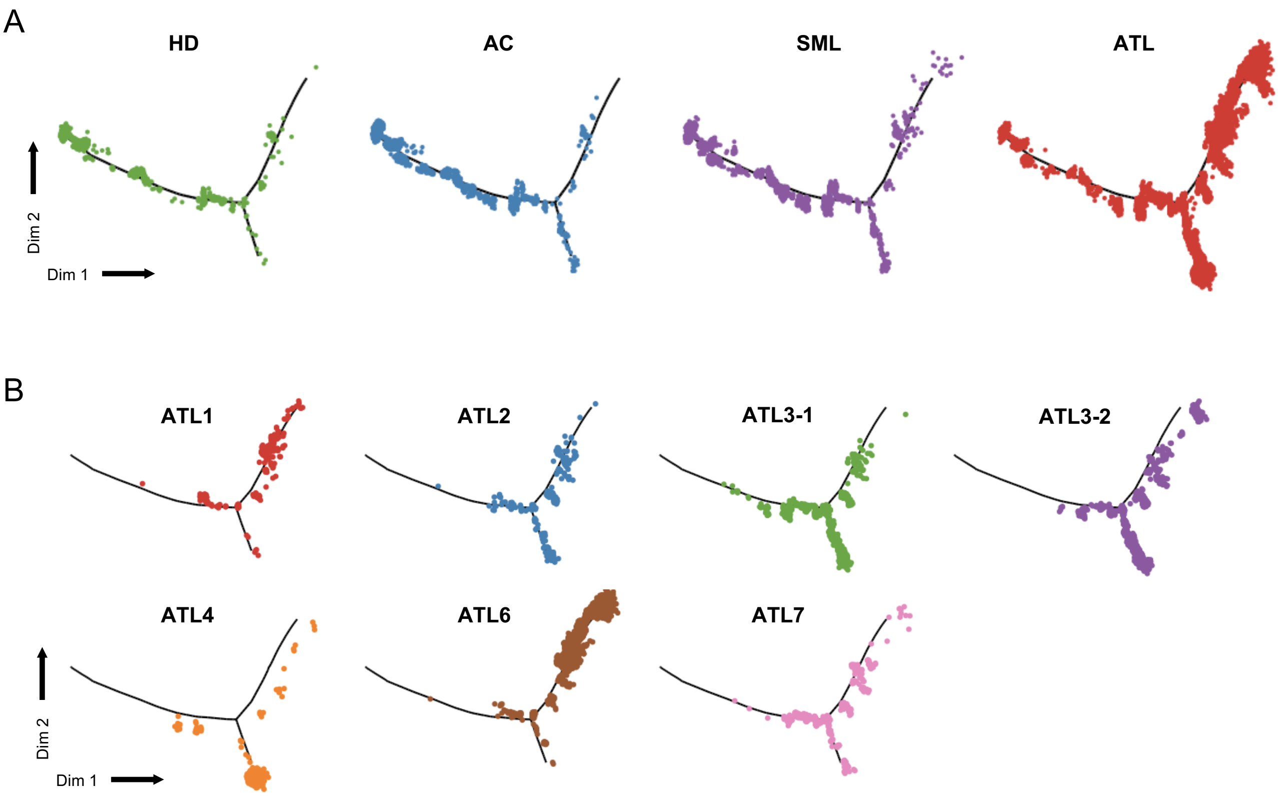


**Figure S7; Pseudotime analysis of all CD4⁺ T-cells.** (**A**) Plots show the distribution of cells along the pseudotime trajectory in Figure **2C** colored by clinical group. (**B**) Plots show the distribution of the most expanded clone for each ATL patient along the pseudotime trajectory.

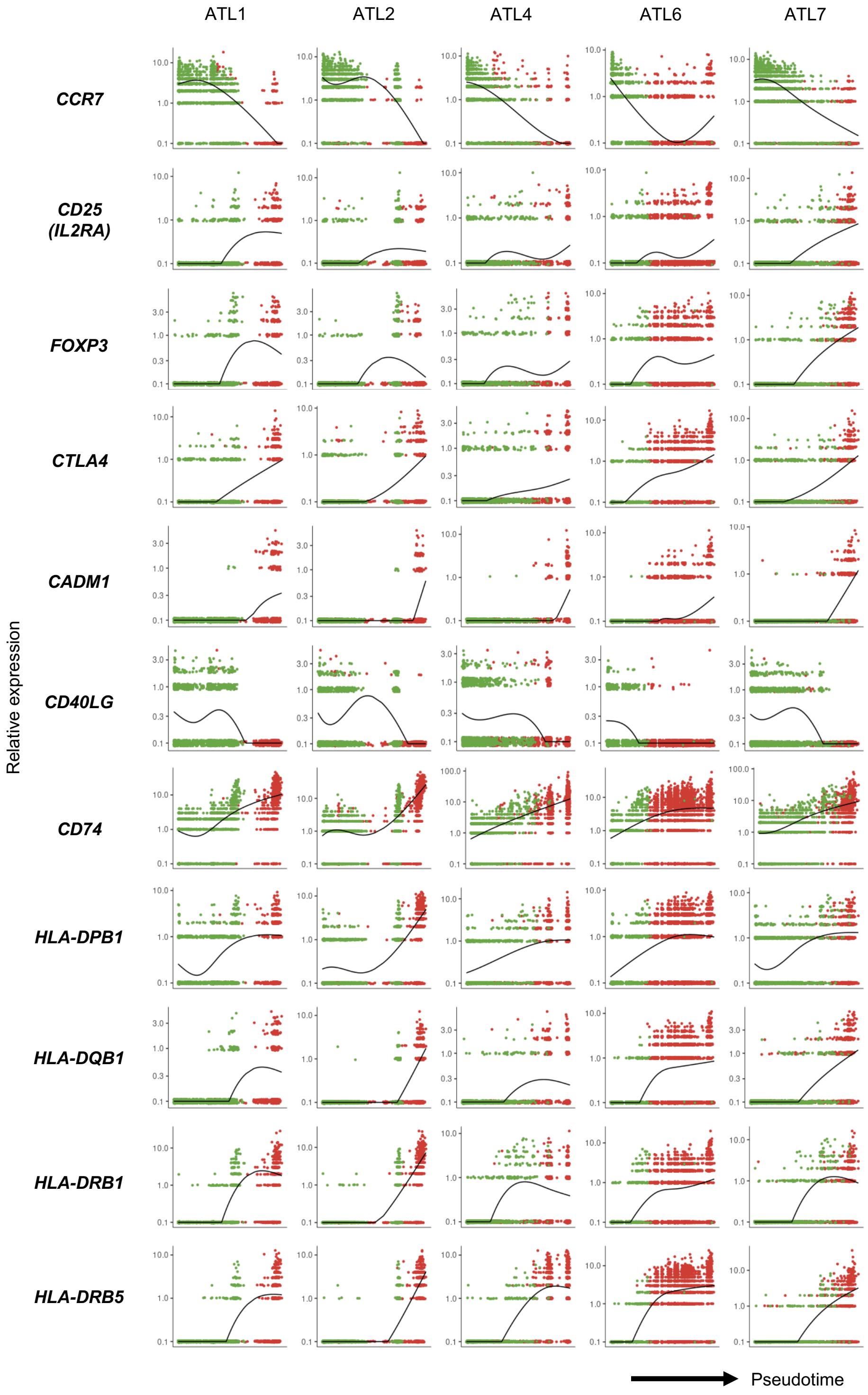


**Figure S8: Pseudotime analysis of single ATL samples.** Expression dynamics of selected T-cell-related and HLA class II genes along the pseudotime axis for individual ATL samples. Color of the dots represent the clinical group as in Figure **3B**.

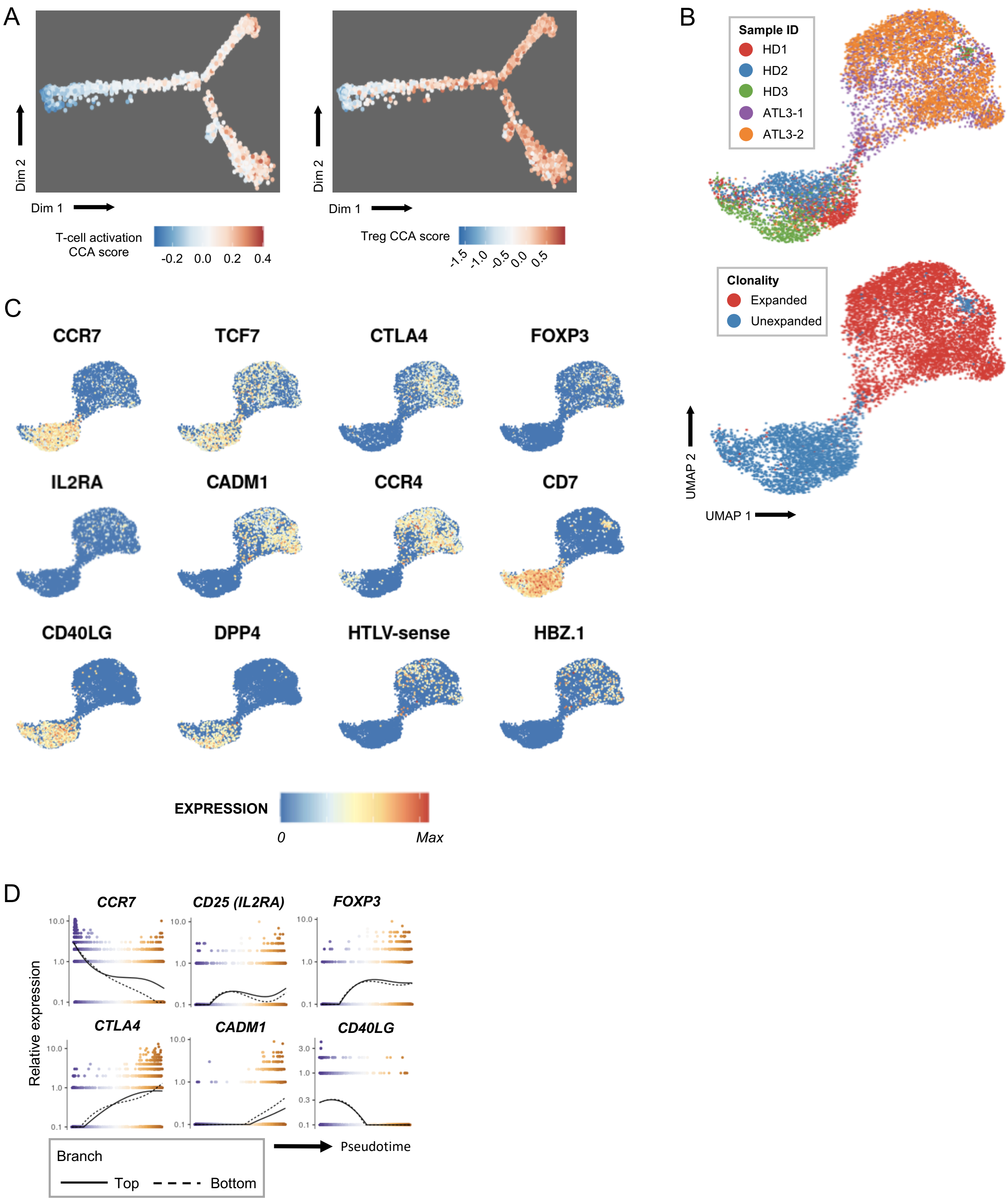


**Figure S9: Clustering analysis of paired ATL sample.** (**A**) Plot showing the distribution of 1-D CCA scores for T-cell activation and Treg in the pseudotime space in Figure **4B**. (**B**) Plots show the clusters identified by Seurat for the 3 HD and 1 paired ATL sample. Left plot is colored by sample ID and the right plot is colored by T-cell clonality. (**C**) Plots show the expression of T-cell and HTLV-1 infection-related genes in the 2-D UMAP space in panel **B**. (**D**) Expression dynamics of selected T-cell-related genes along the pseudotime axis. Color of the dots represent the cells position along the pseudotime axis as in Figure **4B**.

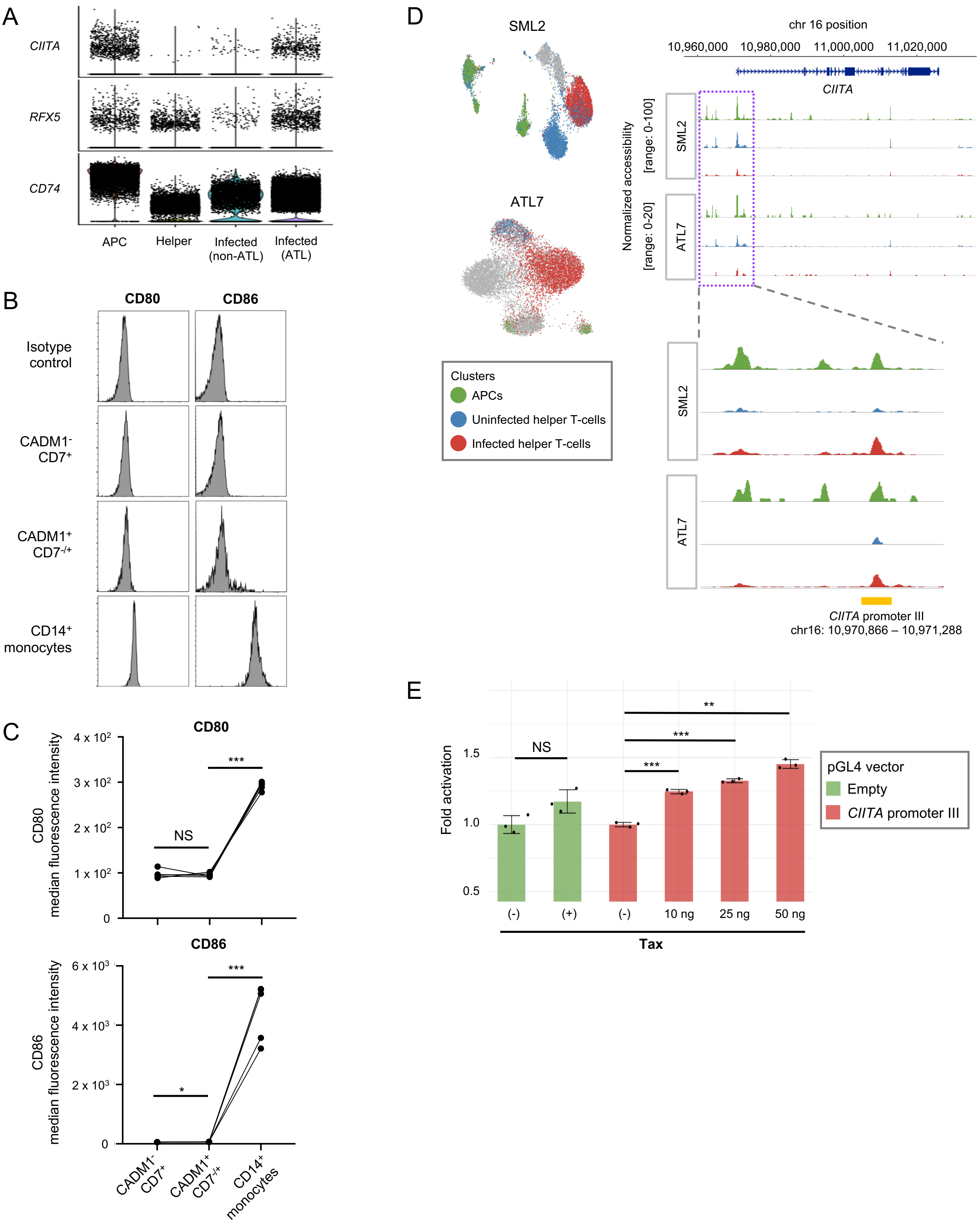


**Figure S10: Expression of HLA class II related genes and CIITA promoter activity.** (**A**) Violin plot shows the expression of HLA class II-related genes in APCs and T-cells. (**B**) Plots show a representative result of CD80 (left column) and CD86 (right column) expression in non-ATL cells (CADM1⁻ CD7⁺), ATL cells (CADM1⁺ CD7⁻/⁺) and monocytes (CD14⁺) from HTLV-1-infected individuals. (**C**) Line graphs show the difference of CD80 (top) and CD86 (bottom) expression between non-ATL cells (CADM1⁻ CD7⁺), ATL cells (CADM1⁺ CD7⁻/⁺) and monocytes (CD14⁺) in 5 HTLV-1-infected individuals. (**D**) Left panel shows the single cell clusters identified from single-cell ATAC-seq (scATAC-seq) of 2 HTLV-1-infected donors. Right panel shows the normalized accessibility peaks in the *CIITA* gene. The color of the peaks corresponds to the cell clusters in the left panel. The zoom-in region shows the peaks around the *CIITA* promoter III. (**E**) Bar graph shows the trans-activation of *CIITA* promoter III with different concentrations of HTLV-1 viral protein Tax. n = 5 (**C**) and 3 (**E**). Data represent mean ± SD. * p-value < 0.05, ** p-value < 0.01, *** p-value < 0.001 by unpaired 2-tailed Student's t-test (**C** and **E**).

**Supplementary Table 1: Clinical data of participants in this study**

| Sample ID | Age | Gender | White blood cell count (/mm$^3$) | Abnormal lymphocyte count (%) | Lymphocyte count (%) | LDH (IU/L) | Serum calcium (mg/dL) | Soluble IL-2 receptor (U/mL) | Proviral load (%) | CADM1$^+$ sub-population on HASFLOW (%) |
|---|---|---|---|---|---|---|---|---|---|---|
| HD1 | 56 | Male | 3,510 | 0 | 34 | - | - | - | - | 1.2 |
| HD2 | 62 | Male | 5,250 | 0 | 36 | - | - | - | - | 1.6 |
| HD3 | 72 | Female | 4,790 | - | - | - | - | - | - | 1.5 |
| AC1 | 61 | Female | 4,790 | 1 | 61 | 170 | 9.5 | 463 | 6.47 | 13.7 |
| AC2 | 69 | Male | 6,360 | 1 | 35 | 167 | 9.4 | 623 | 0.68 | 23.3 |
| AC3 | 76 | Male | 4,080 | 0 | 29 | 209 | 8.8 | - | 0.66 | 17.3 |
| AC4 | 62 | Female | 4,070 | 0 | 44 | 164 | 9.5 | - | Undetectable | 28.8 |
| SML1 | 61 | Female | 3,580 | 4 | 31 | 210 | 9.3 | 666 | 11.91 | 51.2 |
| SML2 | 72 | Female | 7,430 | 8 | 31 | 167 | 9.9 | 529 | 34.60 | 41.9 |
| SML3 | 63 | Female | 4,420 | 2 | 42 | 167 | 9.6 | - | 18.24 | 38.2 |
| ATL1 | 53 | Male | 13,130 | 36 | 16 | 254 | 9.1 | 4,183 | 43.64 | 79.4 |
| ATL2 | 59 | Female | 4,600 | 25 | 18 | 206 | 8.9 | 2,599 | 58.75 | 79.1 |
| ATL3-1 | 71 | Male | 12,590 | 51 | 23 | 267 | 9.5 | 3,981 | 80.63 | 83.1 |
| ATL3-2 | 73 | Male | 13,760 | 53 | 25 | 292 | 9.4 | 4,200 | 66.56 | 80.6 |
| ATL4 | 72 | Male | 18,610 | 10 | 1 | 244 | 10.4 | - | 33.40 | 76.4 |
| ATL6 | 68 | Female | 20,900 | 64 | 12 | 216 | 9.2 | 1,814 | 23.75 | 94.0 |
| ATL7 | 90 | Male | 7,860 | 11 | 22 | 197 | 9.5 | 726 | 40.65 | 80.5 |

**Supplementary Table 2: Primers used for evaluation of T-cell anergy**

| Gene | Forward primer (5' → 3') | Reverse primer (5' → 3') |
|---|---|---|
| *EGR2* | CGAATCCACACTGGGCATAAG | AAACTTTCGGCCACAGTAGTC |
| *EGR3* | AACATCATTAGCCTCATGAGCG | AGGTTGTAGTCAGGAATCATGG |
| *CBL-B* | TACAAGATTCCTTCATCCCACC | TGACCATTATCACAAGACCGAA |
| *DGK-Z* | GAACGACTTCTGTAAGCTCCAG | CTGCTGGTCTGTCTTCATGAG |
| *ITCH* | GCAGCAGTTTAACCAGAGATTC | GTGTGTTGTGGTTGACGAAATA |
| *PDCD1* | AAGGCGCAGATCAAAGAGAGCC | CAACCACCAGGGTTTGGAACTG |
| *HPRT1* | TCGAGATGTGATGAAGGAGATG | CAGCAAAGAATTTATAGCCCCC |
| *IL-10* | GTTGTTAAAGGAGTCCTTGCTG | TTCACAGGGAAGAAATCGATGA |
